# Phagocytosis without mitochondria is a lineage trait of planctomycetal bacteria of prey

**DOI:** 10.64898/2026.10.26.761892

**Authors:** Carmen E. Wurzbacher, Jonathan Hammer, Tom Haufschild, Begüm Serra Büyüktarakçı, Alejandro Jiménez González, Feriel Bouderka, André R. Soares, Julian M. Müller, Luis Felipe Muriel-Millan, María Leticia Ferrelli, Vivien Hotter, Martin Taubert, Kirsten Küsel, Damien Devos, Andreas Klingl, Alexander J. Probst, Siv G. E. Andersson, Nicolai Kallscheuer, Christian Jogler

**Affiliations:** Department of Microbial Interactions, FSU Jena, Germany; Department of Cell and Molecular Biology, Uppsala University, Sweden; Department of Microbial Systems Ecology, Research Center One Health Ruhr of the University Alliance Ruhr, Faculty of Chemistry, University Duisburg Essen, Germany; Plant Development & Electron Microscopy, Biocenter, LMU Munich, Germany; Institut Pasteur de Lille, Univ. Lille, CNRS, Inserm, CHU Lille, U1019 - UMR 9017 - Center for Infection and Immunity of Lille, France; Instituto de Biotecnología y Biología Molecular (IBBM, UNLP-CONICET), Facultad de Ciencias Exactas, Universidad Nacional de La Plata, La Plata 1900, Buenos Aires, Argentina; Department of Aquatic Geomicrobiology, FSU Jena, Germany

## Abstract

Phagocytosis is considered energetically out of reach for prokaryotes. Mitochondria-first models of eukaryogenesis therefore suggest a non-phagocytic archaeal host. Yet two recently described planctomycetes lack mitochondria and engulf entire bacteria. Whether this mode of predation is an oddity of two isolates or the hallmark of a bacterial lineage remained unresolved. Here, we report eight additional predatory *Planctomycetota* from three aquatic habitats. The ten cultured strains now span two classes of the phylum *Planctomycetota*, ‘*Candidatus* Uabimicrobiia’ and one newly proposed class. Using engineered fluorescent prey, we show that nine strains internalize prey into fully sealed compartments whereas one digests its prey externally, a non-engulfing predator for comparison. The strains additionally divide by two distinct modes, however, not correlating with their feeding behavior. Sequence- and structure-based searches detect no eukaryotic phagocytosis protein conserved across the engulfing strains. The actin homologue previously proposed to mediate uptake occurs only in three engulfing strains. Routine engulfment by aerobic bacteria without mitochondria thus unsettles a premise of mitochondria-first models and reopens the question of which cell plans could have hosted the mitochondrial endosymbiont.

## Introduction

Predatory bacteria have been known for more than sixty years^1^. Until recently, four different ways of killing were known: i) from the outside, as epibionts that drain the prey’s cytoplasm (*Vampirococcus*; *Patescibacteriota* - Candidate Phyla Radiation ^2–4^), ii) from within the periplasm (*Bdellovibrio*; *Pseudomonadota* ^5^), iii) by direct invasion of the cytoplasm (*Daptobacter*; phylogenetic affiliation unknown ^2^), and iv) collectively and at a distance, by secreted lytic enzymes (*Myxococcus* ^6^; *Pseudomonadota*). In 2019, Shiratori and colleagues described a fifth mechanism, phagocytosis-like engulfment of prey bacteria and pico-eukaryotic algae (‘*Ca.* Uabimicrobium amorphum’; *Planctomycetota*) ^7^. For organisms displaying this new predatory behavior, we coined the term ‘bacteria of prey’ (BoP) ^8^. Sequence similarity searches against 347 eukaryote signature proteins and the yeast actin-binding proteome returned no homologue of the eukaryotic phagocytosis machinery, apart from a single actin-like protein, suggesting that the uptake is probably analogous rather than homologous to eukaryotic phagocytosis ^7^.

The discovery of a prokaryote that engulfs other cells contradicts a widely held premise of eukaryogenesis: that phagocytosis is energetically out of reach for a cell without mitochondria, so that the alphaproteobacterial endosymbiont must have been acquired via a non-phagocytic mechanism by an archaeal host ^9–11^. Shiratori *et al.* drew inference with care: the organism was not proposed as a descendant of the eukaryotic host, but as a living model in which phagotrophy in the absence of mitochondria could finally be studied ^7^. How much weight that model could bear, however, rested on a single enrichment culture from a single location – and on a phylum with a cautionary history. First described as planktonic fungi, planctomycetes were later recognized as bacteria, then credited with a nucleus-like compartment, endocytosis-like protein uptake and a transitional position on the road to eukaryotes, before each of these eukaryote-like claims collapsed under scrutiny (for review see ^12,13^). We therefore re-sequenced the ‘*Ca.* U. amorphum’ culture to exclude a eukaryotic contaminant, compared the organism with size-matched amoebae, and isolated a second strain, ‘*Ca.* Uabimicrobium helgolandensis’, from the North Sea ^8^. Both strains proved to be unmistakably bacterial and unequivocally distinct from amoebae: they crawl without pseudopodia formation, engulf prey cells they coincidentally touch rather than hunting them at a distance, and divide by a mode found in no other organism, in which the two daughter cells crawl apart on a surface until only a thread-like connection remains, which eventually ruptures due to the physical forces applied. Their genomes lack most canonical cell division and peptidoglycan synthesis genes ^8^.

Environmental sequence data indicated that close relatives to ‘*Ca.* Uabimicrobium’ are widespread ^7^. However, sequence data cannot reveal whether these relatives share the predatory lifestyle and are thus BoP. How widespread phagocytosis-like engulfment is, a peculiarity of a single genus or a trait conserved across a larger phylogenetic breadth, can only be determined from cultures. This phylogenetic question is the first of three that BoP poses. The second concerns the cell biology of engulfment, which challenges long-standing assumptions about the bacterial cell: bacteria are thought to owe their shape to a rigid peptidoglycan (PG) sacculus, a structure that should preclude the envelope deformation required to enclose another cell. Whether engulfed prey ends up fully enclosed within the predator or remains connected to the exterior and what becomes of it thereafter is unknown. The third concerns the molecular basis of uptake and its genomic consequences: the actin-like protein remains the only candidate effector, although membrane coat (MC) proteins, sharing the β-propeller plus α-solenoid architecture of eukaryotic vesicle coats such as clathrin ^14^ were recently reported in ‘*Ca.* U. amorphum’^15^. Whether either protein is conserved across bacteria of prey, as a shared engulfment machinery would require, and to what extent the foreign DNA taken up with every prey cell has shaped the predator’s genome, has not been assessed.

None of these knowledge gaps can be settled by deeper analysis of the two isolates at hand. Comparative genomics requires strains that differ in the trait of interest but are otherwise alike; the two known BoP are indistinguishable in precisely these traits, and no closely related non-engulfing strain exists for comparison. Here, we report eight additional predatory bacteria, obtained in a targeted cultivation campaign, raising the number of cultured predatory *Planctomycetota* to ten. Among them, a single strain feeds externally on its prey rather than engulfing it, providing the natural contrast, engulfing versus non-engulfing, within one phylum, that the questions above require.

## Results

### Previously unknown predatory *Planctomycetota* and their phylogeny

To expand the cultured diversity of predatory *Planctomycetota* beyond the single genus ‘*Ca.* Uabimicrobium’ ^7,8^, we conducted a targeted cultivation campaign, sampling surface water from three sites for predator-selective enrichment: Palau (Pacific Ocean), Heligoland Island (North Sea) and the German Baltic Sea (Fig. 1). Samples from Palau yielded ‘*Ca.* Uabimicrobium palauense’ PEnr_61_6, a new species within the genus ‘*Ca*. Uabimicrobium’. Heligoland Island samples yielded two strains of a new genus that belong to the same species, ‘*Ca.* Ligurimicrobium helgolandense’ HIEnr_11 and HIEnr_12, and one strain representing a new class within the phylum *Planctomycetota*, ‘*Ca.* Nanovorax helgolandensis’ HIEnr_7_s (GTDB class c J058, SILVA clade BD7-11). The Baltic Sea samples yielded four strains forming the new genus, comprising two species represented by strains BSEnr_70 as well as BSEnr_200, BSEnr_86_b and BSEnr_24_14, respectively (see Supplementary Table 1 – 8 for details). All strains are maintained as enrichment cultures with *Alteromonas macleodii* as prey while none of them are monoxenic.

**Fig. 1.**
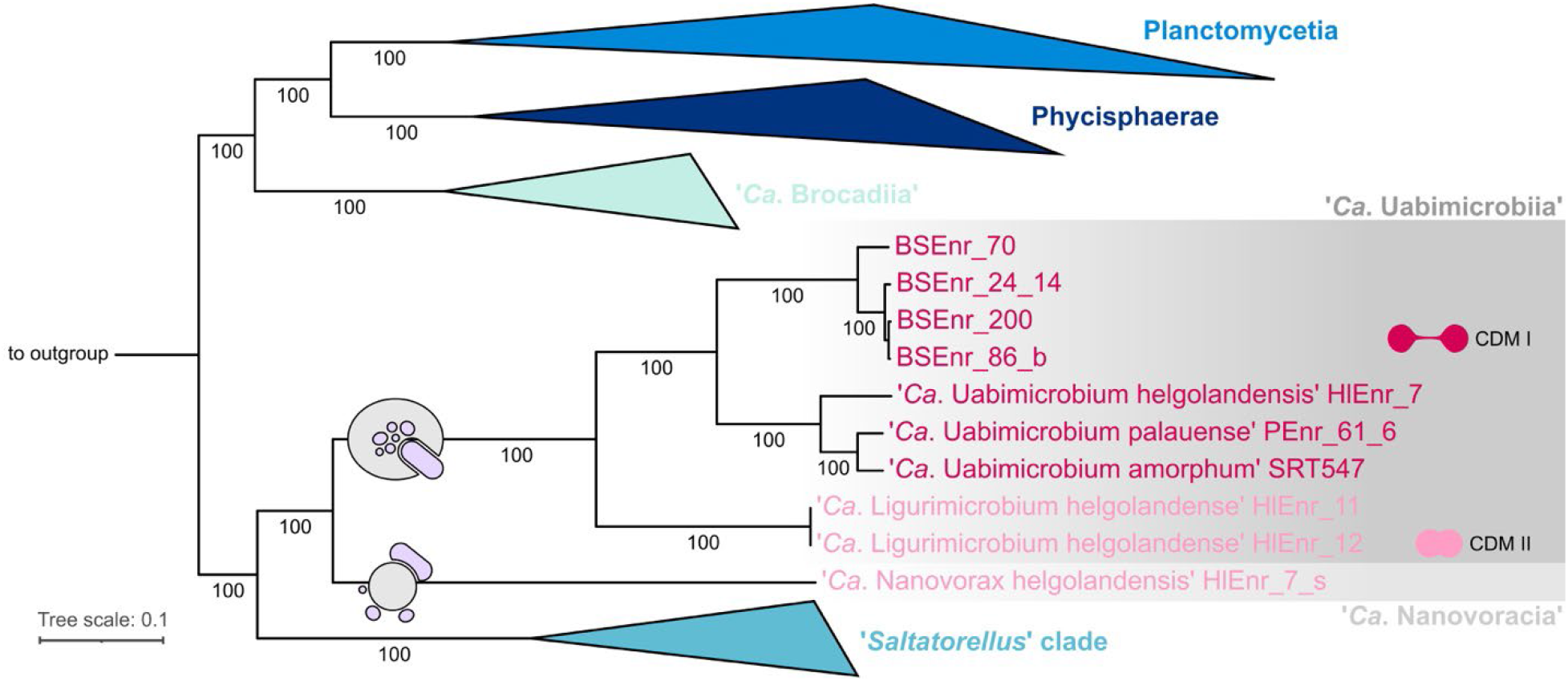
Whole genome-based phylogeny with DNAS G+C content correction of the new strains. The classes ‘*Ca*. Uabimicrobiia’ (engulfing, bacteria of prey) and ‘*Ca*. Nanovoracia’ class. nov. (external digestion) form a monophyletic sister clade to the ‘*Saltatorellus*’ clade. While cell division mode I (CDM I) can only be found in members of ‘*Ca*. Uabimicrobiia’, members of both classes seem to exhibit cell division mode II (CDM II).

Reconstructing a robust phylogeny on class-level of the phylum *Planctomycetota* to determine the cultured predatory bacteria position within the phylum is impeded by substantial variation in the genomic DNA G+C content^16^. Across the phylum, DNA G+C contents range from ∼34–39% in the predatory group to ∼70% in the ‘*Saltatorellus*’ clade — deviations large enough to compromise clustering and mask phylogenetic signal ^17^. Thus, we corrected for DNA G+C content differences in our phylogenetic reconstruction as previously described ^16^. Our phylogenetic tree (Fig. 1) shows the two classes of predatory *Planctomycetota*, the described class ‘*Ca.* Uabimicrobiia’ and the previously uncultured class ‘*Ca.* Nanovoracia’, to form a sister clade to the ‘*Saltatorellus’* clade. The three classes *Planctomycetia*, *Phycisphaerae* and ‘*Ca.* Brocadiia’ are in turn forming a monophyletic sister clade to the ‘*Saltatorellus*’, ‘*Ca.* Uabimicrobium’ and ‘*Ca*. Nanovoracia’ clades. Metagenome-assembled genomes (MAGs) suggest members of the previously unknown class ‘*Ca*. Nanovoracia’ (SILVA r144 BD7-11 clade, GTDB c J058) to be present in various environments, including groundwater (Supplementary Fig. 1). Thirteen MAGs cluster with the ‘*Ca.* Uabimicrobiia’ clade (SILVA r144 Candidatus Uabimicrobiia, formerly Pla4 lineage, GTDB c Uabimicrobiia), mostly associated with wastewater and sludge (Supplementary Fig. 1).

### Nine of ten predatory *Planctomycetota* engulf their prey

To determine whether the previously unknown strains engulf their prey completely, we first developed a protocol to genetically manipulate the prey bacterium *Alteromonas macleodii* (see Material and methods). We then constructed two fluorescent strains of *A. macleodii* prey bacteria, which constitutively express genes either encoding GFPmut2 or TurboRFP in the cytoplasm. Time-lapse microscopy revealed uptake of these mutant prey bacteria by all members of the class ‘*Ca.* Uabimicrobiia’ (Fig. 2 a). By contrast the only member of the second class, ‘*Ca.* Nanovorax helgolandensis’ HIEnr_7_s, does not engulf its prey (Fig. 2 b). No prey-derived fluorescence was observed within its cells, and prey cell leftovers were frequently visible in SEM micrographs (Fig. 2 c). Instead, the strain degrades prey on the outside of its cell. After reversible attachment, it appears to alter the prey envelope locally and, like its engulfing relatives, dissects the prey into smaller bounded bodies, necessarily bounded by the prey’s own envelope, since the prey is never located inside the predator. As in engulfing BoP, fluorescence is retained in bodies of every size. ‘*Ca.* N. helgolandensis’ HIEnr_7_s thus preys on the same organisms as its phagocytic relatives without performing bacterial phagocytosis. It provides the contrast that earlier comparative analyses lacked: subtracting its gene content from that of the phagocytic strains should narrow the list of candidate genes for bacterial phagocytosis substantially.

**Fig. 2.**
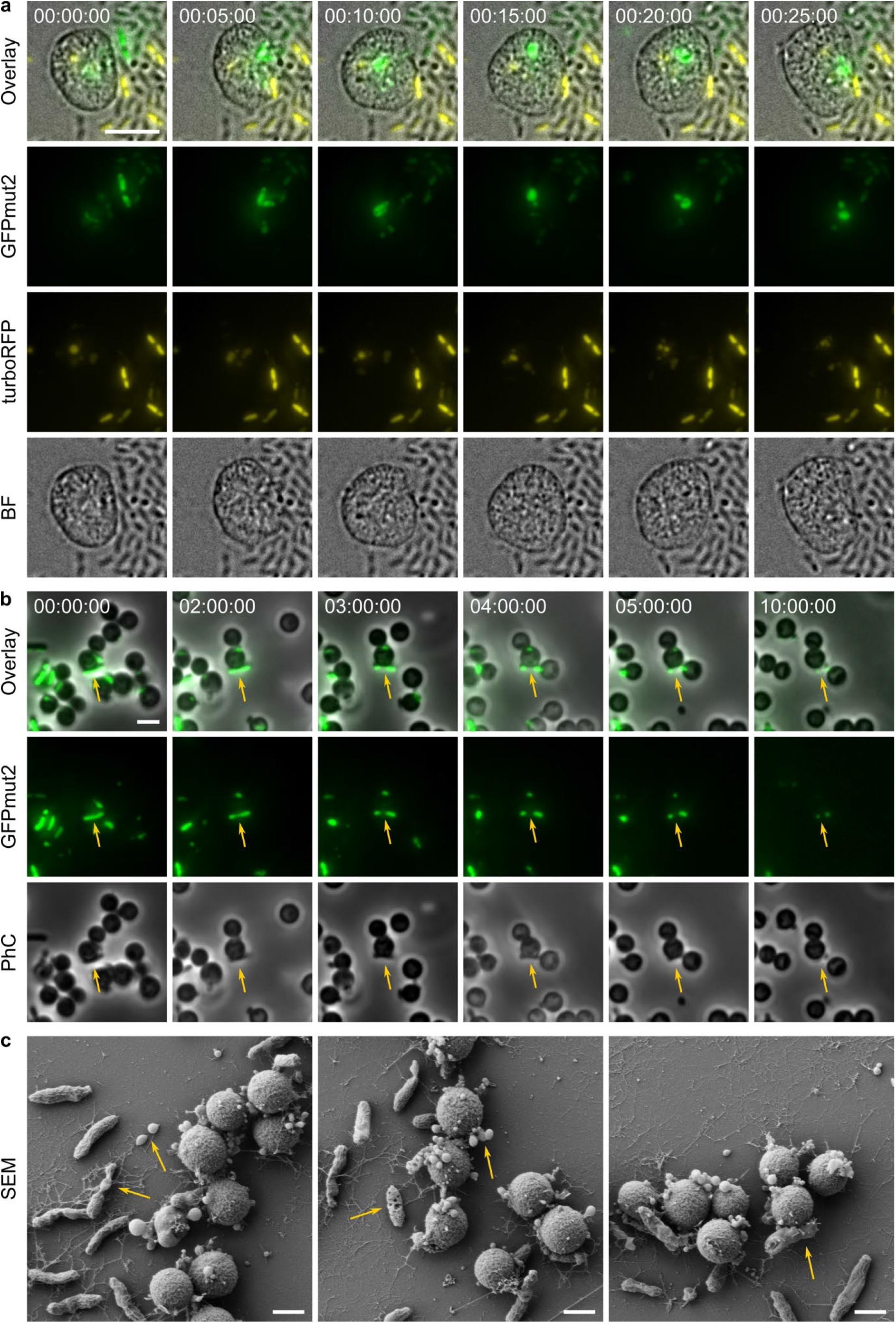
Bacterial phagocytosis by ‘*Ca.* Uabimicrobium amorphum’ SRT547 contrasted with external digestion by ‘*Ca.* Nanovorax helgolandensis’ HIEnr_7_s. (a) Time-lapse fluorescence microscopy of ‘*Ca.* U. amorphum’ SRT547 preying on *Alteromonas macleodii* expressing genes encoding GFPmut2 (green) or TurboRFP (yellow). Engulfed prey cells rapidly lose their shape and become intracellular fluorescent bodies; a derived body resulting from an earlier uptake event (yellow) is split into smaller fluorescent bodies during the same period. Merging of differently coloured bodies was not observed. (b) ‘*Ca.* N. helgolandensis’ HIEnr_7_s does not engulf prey cells; prey is instead digested externally and similarly split into smaller bodies. (c) scanning electron microscopy of HIEnr_7_s, showing severe damage to prey cells and numerous vesicle-like structures — likely prey undergoing digestion — on the predator’s surface. Scale bars are 5 µm (a), 2 µm (b) and 1 µm (c). Timestamps, hh:mm:ss.

### Bacteria of prey engulf their victims in fully sealed compartments

Whether engulfment by BoP creates a compartment that is closed off from the exterior has remained an open question. Shiratori *et al.* observed narrow ducts connecting some prey-containing vacuoles (phagosome-like vacuoles, PLVs) to the cell exterior or to other PLVs ^7^. We therefore asked whether the compartment is sealed and what happens to the prey within it. Our fluorescent *A. macleodii* prey strains allowed the fate of ingested cells to be followed over time (Fig. 2 a). After uptake, prey cells remain fluorescent and initially keep their rod shape. Over 10-15 minutes the signal becomes round, indicating loss of the prey’s cellular integrity, but it does not disperse into the predator’s cell; the prey-derived material remains confined. The confined signal then fragments into several smaller fluorescent bodies, and fluorescence is retained in bodies of every size, so each must be bound by a barrier confining the soluble cytoplasmic protein — presumably a membrane. Bodies derived from GFP- and RFP-labelled prey never merged in any of our recordings, thus, material from different prey cells is kept separate throughout digestion.

Fragmentation of single prey cells into several bounded bodies implies that membranes are remodeled during digestion. Which membrane forms the boundary — the prey’s own envelope, the membrane of the predator-derived compartment that Shiratori *et al.* described as the phagosome-like vacuole (PLV) ^7^, or both — is beyond the resolution of fluorescence microscopy. We therefore refer to the compartment as the PLV and to the fluorescent units simply as prey-derived bodies. The same limitation applies to the question of whether PLVs are sealed. To resolve it, we imaged ‘*Ca.* Uabimicrobium amorphum’ by focused-ion-beam milling and scanning electron microscopy (FIB/SEM) and reconstructed cell volumes in three dimensions (Fig. 3 a, Supplementary Movie 1). PLVs containing prey (Fig. 3a, red) proved to be fully enclosed, with no connection to the exterior. During reconstruction, we found BoP DNA to be organized in partly separated compartments as well (Fig. 3 a, blue). Transmission electron microscopy (TEM) furthermore revealed that the membrane situation around PLVs is exactly opposite to the envelope of the predator (Fig. 3 b). Prey is thus taken up into a closed compartment. TEM thin sections of BoP cells showed a largely empty cell body that, besides enclosed PLVs, contained interconnected cytoplasmic compartments (Fig. 3 b). SEM further revealed that engulfment of rod-shaped prey was initiated either at a prey’s cell pole (polar engulfment) or along its lateral surface (lateral engulfment) (Fig. 3 c). During engulfment, an opening of the predator’s envelope occasionally remained visible after the prey cell had been internalized potentially leading to the ducts observed by Shiratori *et al.* in TEM thin sections (Fig. 3 c). In some cases, prey cells appeared to be folded in half during lateral engulfment, before internalization (Fig. 2 a, green cell and Fig. 3 c).

**Fig. 3.**
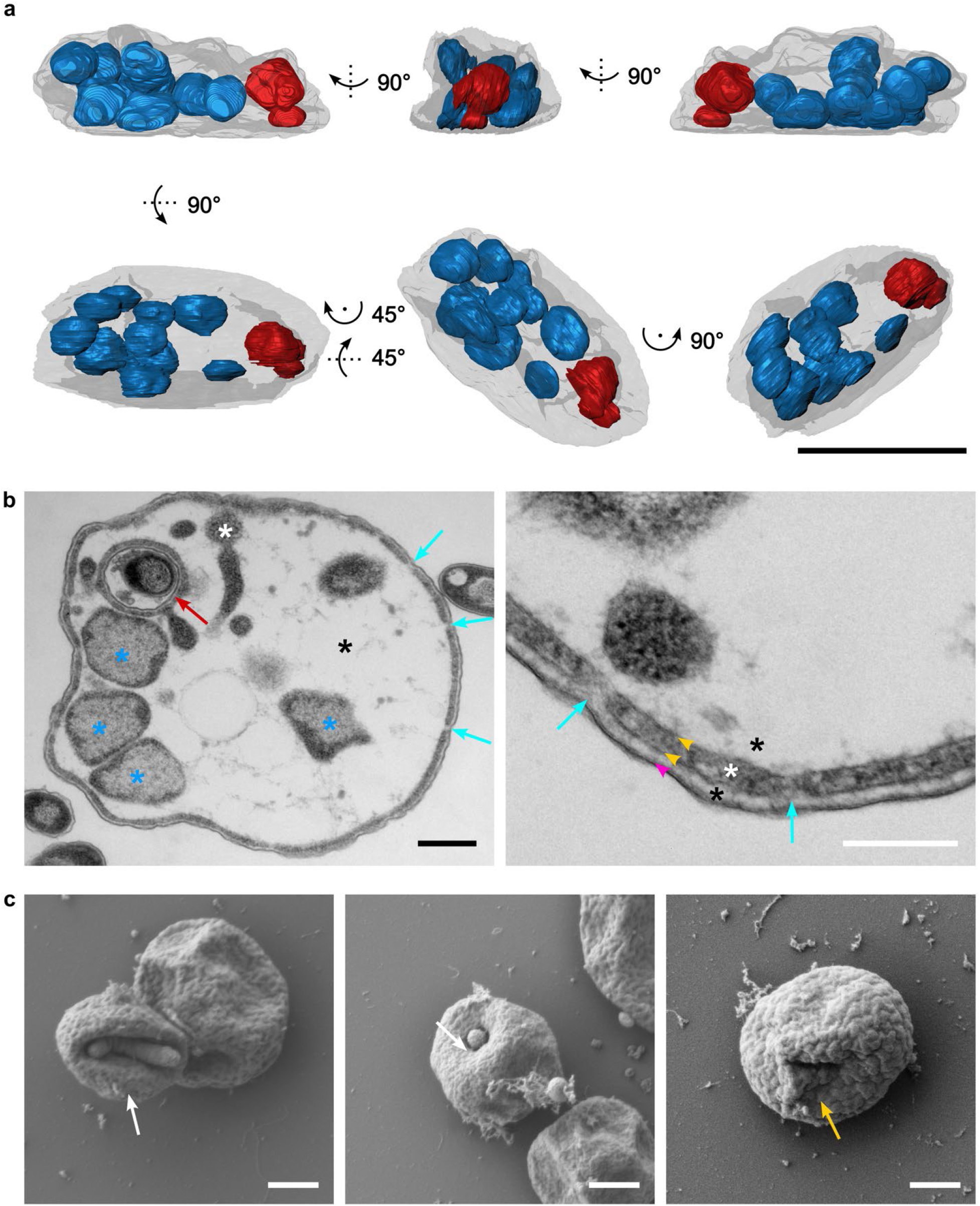
Electron microscopy of ‘Ca. U. amorphum’. (a) FIB-SEM 3D reconstruction of DNA (blue) and phagosome-like vacuoles (red). The vacuoles are fully enclosed compartments inside the cell without any remaining connection to the exterior. (b) TEM thin sections. A thin layer of cytoplasm, enclosed by a symmetric lipid bilayer (yellow arrowheads), lines the cell periphery and surrounds phagosome-like vacuoles (red arrow), from which invaginations extend into the cell body (white asterisks). The peripheral cytoplasm shows discontinuities (cyan arrows) that connect the space beneath the outer, asymmetric lipid bilayer (magenta arrowhead) with the cell interior, suggesting that most of the cell volume is periplasm (black asterisks). Blue asterisks, DNA. (c) SEM micrographs of prey engulfment showing lateral (left) and polar (middle) engulfment of prey cells (white arrows). Some cells display openings in the cell surface (right, yellow arrow), probably representing a late stage of engulfment. Scale bars, 5 µm (a), 500 nm (b, left), 200 nm (b, right), 1 µm (c).

### No eukaryotic phagocytosis machinery is conserved across engulfing strains

To test whether bacterial phagocytosis shares molecular components with eukaryotic phagocytosis, we compiled a curated list of 192 eukaryotic phagocytosis proteins (Supplementary Table 9) and searched all BoP genomes for sequence homologs (threshold E < 10^-4^, Supplementary Table 10). In parallel, we predicted the protein structures and performed a structure-based annotation, because homology between bacterial and eukaryotic proteins can persist in structure after sequence similarity has decayed beyond detection. At the permissive threshold chosen to maximize sensitivity (E < 10⁻⁴), the profile searches additionally returned distant, mostly partial hits to ancient protein superfamilies represented in the phagosome proteome, including actin-superfamily proteins and AAA+ ATPases related to NSF and VCP (Supplementary Table 10). None of these hits is specific to the engulfing strains, and structure predictions of representative hits did not support the eukaryotic assignments. Neither approach thus recovered a eukaryotic phagocytosis protein specifically conserved among BoP.

Yet, two previously known candidates required closer inspection. Membrane coat (MC) proteins — clathrin, the COPI and COPII vesicle coats as well as the scaffold nucleoporins — share an architecture of β-propeller domains and α-solenoids ^18^. Bacterial MC-like (MCL) proteins with this architecture have been reported from *Planctomycetota* ^14,15,19^. Using bacterial MCL proteins and profile HMMs of six yeast MC proteins as queries, we identified 12 MCL proteins in the new isolates (Fig. 4). Each of the eight new phagocytic strains encodes a homolog of *Gemmata obscuriglobus* MCL18 (Fig. 4 a), as does ‘*Ca.* U. amorphum’ ^15^. The same homolog was, however, also detected in the non-engulfing ‘*Ca.* N. helgolandensis’ HlEnr_7_s. Four of the phagocytic strains (PEnr_61_6, HlEnr_7, HlEnr_11 and HlEnr_12) additionally encode a homolog of *G. obscuriglobus* MCL19, which also matches the COPII outer-coat protein Sec31 (Fig. 4 b). An MCL19 homolog has previously been reported from ‘*Ca.* U. amorphum’ ^15^. MCL proteins are thus conserved among the BoP but not restricted to them, as they are found in other *Planctomycetota*, and some relatives ^15,20^.

**Fig. 4.**
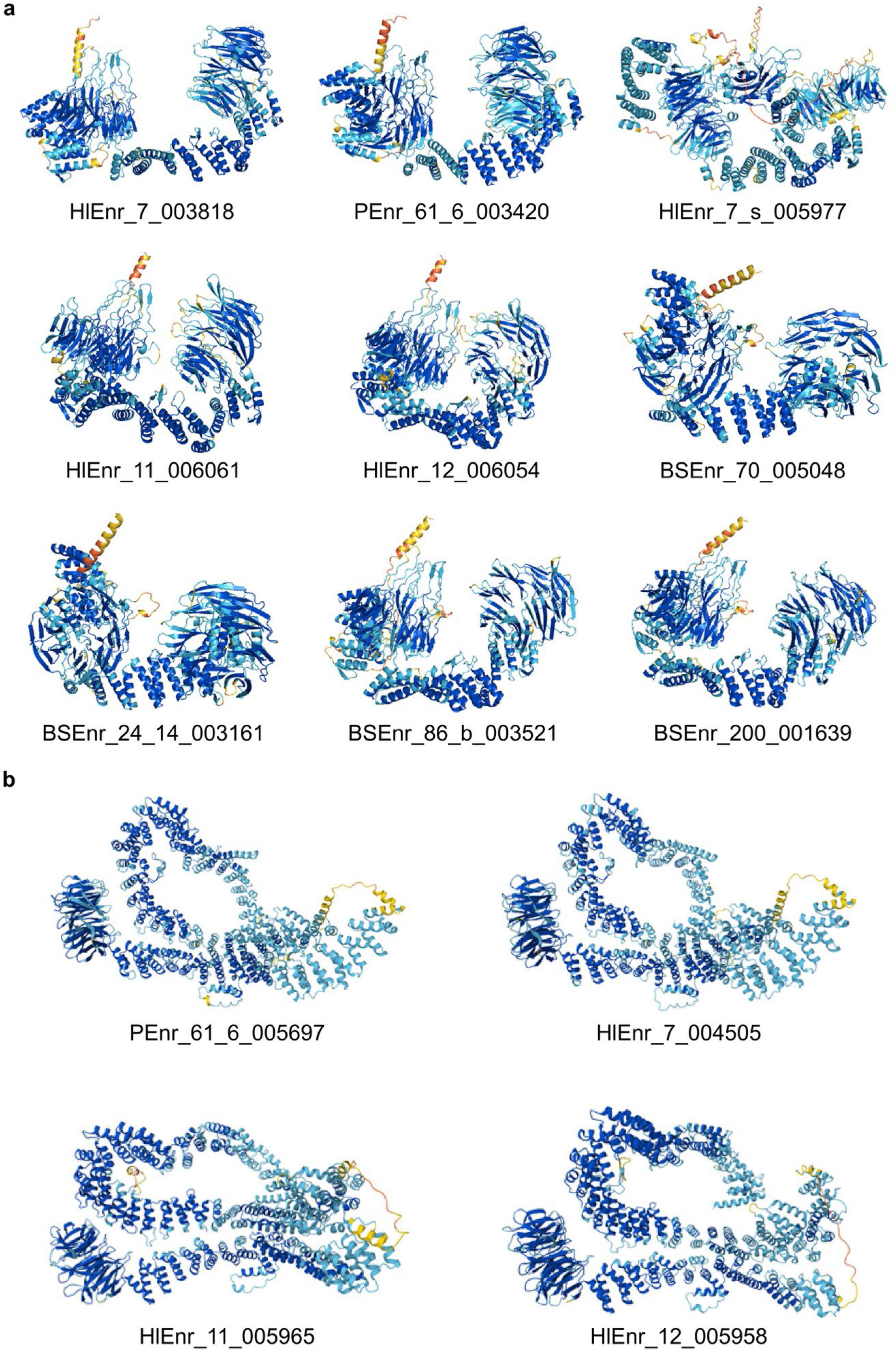
Bacteria of prey encode eukaryotic membrane coat-like proteins. Predicted structures of the 12 membrane coat-like (MCL) proteins found in the proteomes of the new isolates. They were found by BLASTp searches with bacterial MCL proteins and by profile-HMM searches (jackhmmer/hmmsearch) with *Saccharomyces cerevisiae* membrane coat (MC) proteins as queries. (a) Eight MCL proteins homologous to *G. obscuriglobus* MCL18, one in each of the eight engulfing strains, made up of two β-propeller domains and an α-solenoid segment. Proteins in the same row of a and the first column of b come from the same strain. No MCL protein was detected in the non-engulfing strain ‘*Ca.* Nanovorax helgolandensis’ HlEnr_7_s. Models were predicted with ColabFold or AlphaFold 3 and are colored by per-residue confidence (pLDDT): dark blue, >90; light blue, 70–90; yellow, 50–70; orange, <50. Labels are locus tags, and the prefix identifies the strain. (b) Four MCL proteins homologous to both MCL19 of *Gemmata obscuriglobus* and the COPII outer-coat protein Sec31 of *S. cerevisiae*, one each in strains PEnr_61_6, HlEnr_7, HlEnr_11 and HlEnr_12. Each combines a β-propeller domain with an extended α-solenoid. MC, membrane coat; MCL, membrane coat-like.

The second candidate was a lokiactin homolog, the first protein proposed to mediate bacterial phagocytosis in ‘*Ca.* U. amorphum’ ^7^. Homologs were found in only two of the eight new phagocytic strains (PEnr_61_6 and BSEnr_200) and were entirely absent from five genomes. The genome of BSEnr_86_b, however, encodes 93 amino acids of the N-terminus of the BSEnr_200 homolog in the same genomic context annotated as a hypothetical protein. The lokiactin homolog is therefore not conserved across the BoP. In summary, no protein in the dataset was both conserved across all nine BoP and absent from the non-engulfing strain.

Since our targeted approach did not return any conserved eukaryotic phagocytosis-related proteins, we performed an untargeted comparative genomics approach, computing the pan-genome of all BoP and subtracting gene families that were found in the genomes of all other *Planctomycetota* members: 245 orthogroups were found exclusively in BoP, including new types of serine/threonine kinases, which is consistent with a dramatic expansion of this protein family in prokaryotic and eukaryotic cells with phagocytosis-like behaviors ^16^ (Supplementary Table 11). None of them is homologous to a known phagocytosis protein, and no individual protein stands out as a key effector. Whether any of them participates in uptake will require mutant generation in a BoP, which is not yet possible.

By further analyzing the genomes of predatory *Planctomyetota*, it became evident that their gene content, in contrast to canonical model *Planctomycetota* such as *Planctopirus limnophila*, *Gemmata obscuriglobus*, *Stieleria maiorica* and *Isosphaera pallida* points towards markedly elevated levels of horizontal gene transfer (HGT) (Supplementary Fig. 2). In an eggNOG-mapper best-hit analysis, only an average of 13.5% genes were assigned to *Planctomycetota*. The remainder derived predominantly from other bacterial phyla (on average 34 per genome), with a small fraction assigned to eukaryotes, including *Amoebozoa*. A complementary BLASTp-based assignment of the model strain genomes confirmed that the overwhelming majority (>90%) of the gene content is of planctomycetal or other bacterial origin, with only marginal cross-domain input (Supplementary Fig. 3).

### Feeding mode and cell division mode are decoupled

The previously described BoP divide by a mode that requires surface-associated motility (Cell Division Mode I; CDM I): the two poles of the mother cell migrate in opposite directions stretching the middle of the cell body to a thread-like connection up to three cell diameters long, which finally ruptures (Fig. 5 a) ^7,8^.

**Fig. 5.**
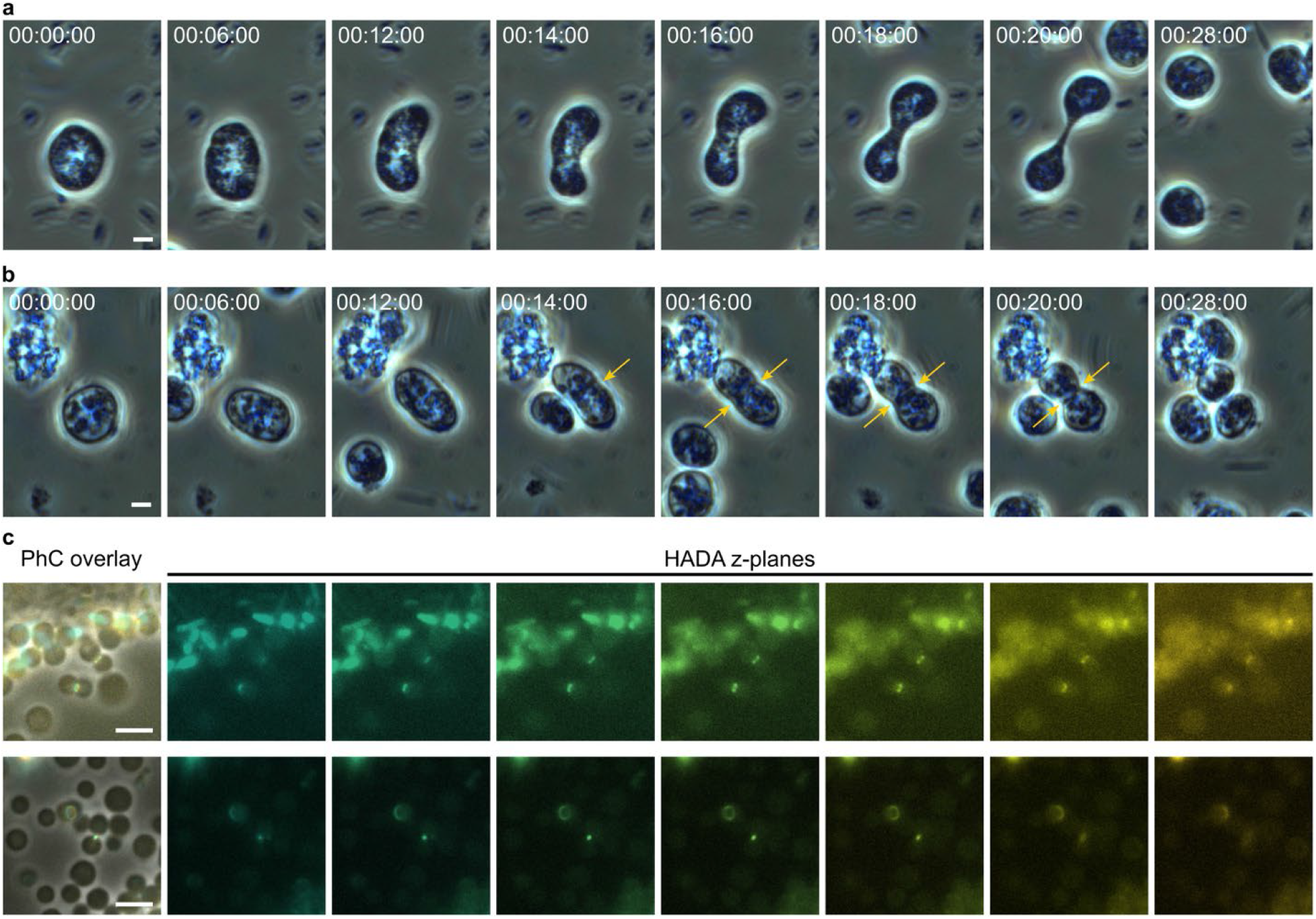
Cell division of predatory Planctomycetota. (a) Most strains employ the hitherto described mode of cell division among predatory *Planctomycetota*, which is characterized by the two emerging daughter cells walking apart, leading to thinning and extension in the middle of the cell until only a thread like connection is left that eventually disrupts.) (b) Some strains employ a previously unknown mode of cell division (CDM II), which strikingly resembles binary fission. Yellow arrows indicate the site of constriction. (c) In the class ‘*Ca.* Nanovoracia’ class. nov., the previously unknown mode of division is exhibited through a peptidoglycan ring at the constriction site. Peptidoglycan was visualized using HADA, z-stacks were recorded and color coded. The resulting composite image of all z-planes is displayed at the left together with a phase contrast (PhC) overlay. Scale bars are 2 µm, timestamps are in hh:mm:ss format.

Two of the novel BoP, however, were found to divide differently. The two ‘Ca. Ligurimicrobium helgolandense’ strains HIEnr_11 and HIEnr_12 constrict at mid-cell in a manner resembling binary fission (CDM II; Fig. 5 b). The same is also true for the non-engulfing ‘Ca. Nanovorax helgolandensis’ HIEnr_7_s. Thus, feeding mode and division mode are decoupled across the strain set, which allowed us to compare the genomes by division mode independent of bacterial phagocytosis.

A presence/absence comparison of orthogroups between strains dividing by CDM I and CDM II identified 478 orthogroups that differed between the two division modes (Supplementary Table 12). Of these, 344 were present in CDM I strains but absent from CDM II strains, and 134 showed the reverse pattern. The latter group included a penicillin-binding protein (PBP), whose gene is located downstream of *mreD* in all CDM II genomes. Closer inspection revealed that CDM I strains encode a truncated version of this protein, initially annotated as hypothetical, in the same genomic context (Fig. 6 a, Supplementary Fig. 4). CDM II strains additionally encoded more genes associated with PG synthesis and remodeling than CDM I strains (Fig. 6 b). Differences in peptidoglycan metabolism may therefore distinguish the two division modes.

**Fig. 6:**
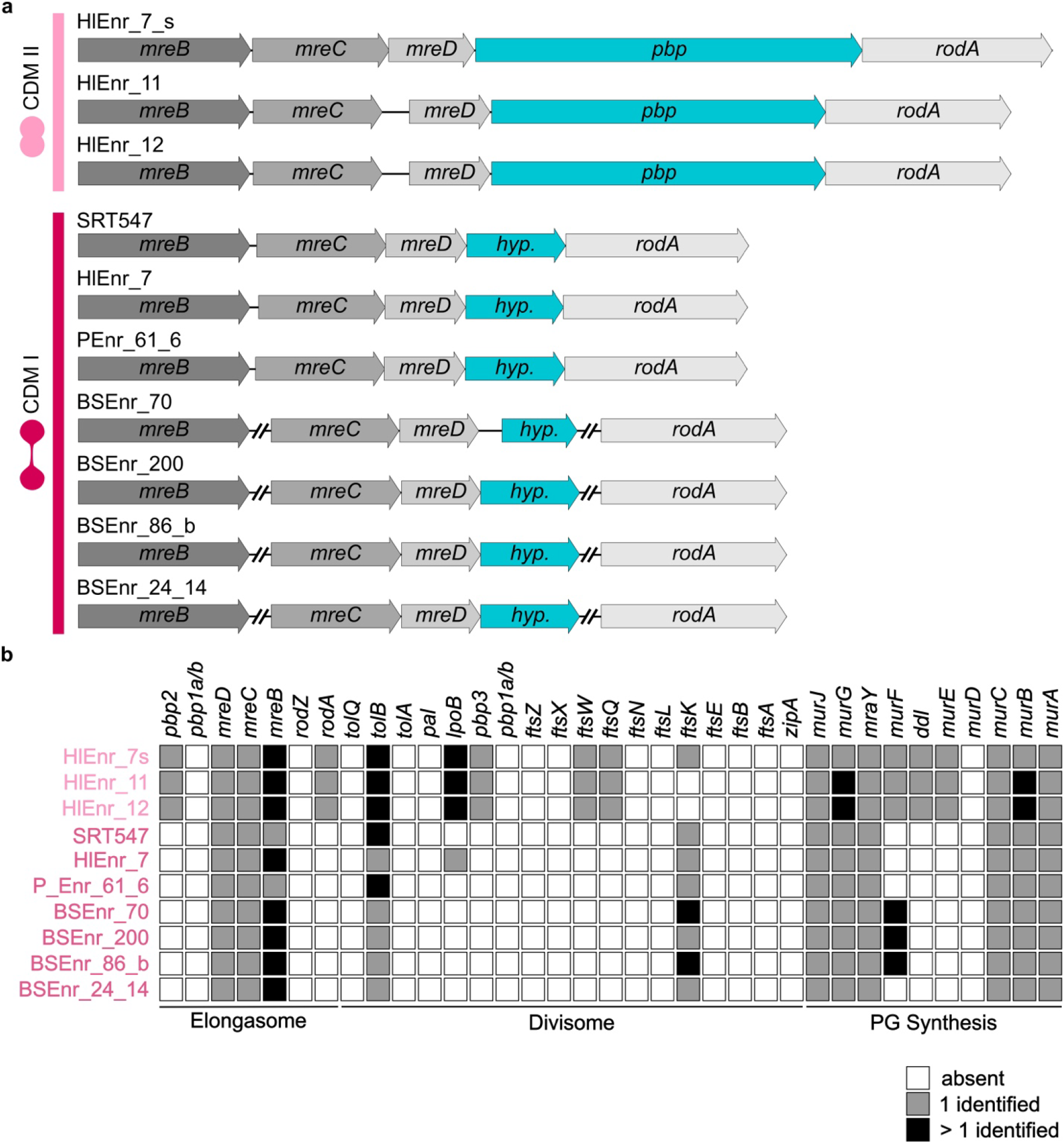
Genomic context of peptidoglycan synthesis and cell division related genes. (a) Elongasome genes in predatory *Planctomycetota* members. In CDM II strains, a penicillin-binding protein (*pbp*) follows *mreD*, while in CDM I strains, only a truncated version (*hyp*.) is found, which is annotated as “hypothetical protein”. (b) Gene presence/absence analysis based on KOfams. KOfams absent in the genome are indicated by a white square, only one matching KOfam is displayed with a grey square and more than one found KOfam is illustrated by a black square. Predatory *Planctomycetota* share many, but not all, genes for peptidoglycan peptidoglycan synthesis while most KOfams representing genes of the divisome could not be identified. Of the elongasome, the mre system could be identified in all Predatory *Planctomycetota*. Based on this bioinformatical analysis, both cell division modes could be differentiated: genomes of organisms performing CDM II contained additional genes for PG synthesis (most importantly *ddl*) as well as genes coding for divisome and elongasome components, most notably *ftsQ*, *ftsW*, *pbp3* and *rodA* and *pbp2*. While yet to be confirmed *in vivo*, the analysis supports the differentiation between both cell division modes and identifies potential candidates accounting for the differences in BoP cell division.

To test this *in vivo*, we labelled nascent PG in all ten strains with the fluorescent D-amino acid HADA, which shows sites of active PG incorporation. Only in ‘*Ca*. N. helgolandensis’ HlEnr_7_s a signal in the division plane could be observed: a highly dynamic ring structure, lost in living cells around 8 min after starting the first washing step (Fig. 5 c). Yet, no signal consistent with a cell-spanning sacculus was observed. Strains belonging to the class *Ca*. Uabimicrobiia, by contrast, only showed signals likely resulting from uptake of the labelled amino acid during feeding. Because the specific signals in ‘*Ca.* N. helgolandensis’ HIEnr_7_s were rather faint, a signal of a comparable strength would be difficult to detect in members of the class ‘*Ca*. Uabimicrobiia’ against this background. The only distinct signals inside their cells were occasional prey-shaped structures, which most likely derive from the prey’s own PG, since prey cells not yet engulfed showed HADA labelling throughout their envelope. Thus, we neither detected a sacculus nor a division plane ring in ‘*Ca*. Uabimicrobiia’ but cannot conclude from labelling alone whether these strains synthesize PG. The genomic difference identified by the subtractive comparison could thus only be confirmed *in vivo* for the one strain in which peptidoglycan synthesis is detectable.

## Discussion

BoP, previously known from two strains of a single genus, now comprise nine strains in three genera of ‘*Ca.* Uabimicrobiia’ ^7,8^. The eighth of our previously unknown isolates, ‘*Ca.* Nanovorax helgolandensis’ HIEnr_7_s, is no BoP by our definition. It preys on the same bacteria as its phagocytic relatives but digests them externally, rather resembling epibiotic predation — though unlike *Vampirococcus*, the predator is far larger than its prey^2^. To our knowledge, neither epibiotic predation within the *Planctomycetota* nor an epibiotic predator that dissects its prey has been described before. As a predator that does not perform bacterial phagocytosis, ‘*Ca.* N. helgolandensis’ HIEnr_7_s supplied the contrast earlier comparative analyses lacked, and subtracting its gene content from the BoP pan-genome narrowed the candidate list to 245 orthogroups. Whether bacterial phagocytosis occurs elsewhere in ‘*Ca.* Nanovoracia’ remains unknown: HlEnr_7_s is the only cultured member of the class, and the metagenome-assembled genomes available for comparison (Supplementary Fig. 1) do not reveal the feeding mode.

A second contrast for comparative analyses runs orthogonally to the first: Strain HIEnr_7_s divides by a binary-fission-like mode (CDM II), as do the phagocytic strains HIEnr_11 and HIEnr_12, whereas the remaining BoP divide as previously described (CDM I) ^8^. Feeding mode and division mode are therefore decoupled across our strain set, and the two traits could be analysed independently. Thus, CDM I is not universal among all BoP. For division, the comparison implicated PG as a key difference: CDM II strains encode a full-length penicillin-binding protein between the *mreBCD* gene cluster and *rodA* that is truncated in CDM I strains, alongside an expanded complement of PG synthesis and remodelling genes. Finally, HADA labelling revealed a transient, highly dynamic ring confined to the division plane of HIEnr_7_s while no signal consistent with a sacculus was observed. The exact role of this ring in cell division is unknown. The arrangement is reminiscent of the *Chlamydiota*, long held to lack PG entirely until D-amino acid probes revealed pathogenic *Chlamydia* to assemble a narrow, transient ring in mid-cell instead of building classical PG sacculi ^21–23^. HADA labelling was also applied to ‘*Ca*. Uabimicrobiia’ strains of both division modes, but only HIEnr_7_s, the non-engulfing CDM II strain, gave a conclusive signal. The absence of clear signals for either a division ring or a sacculus in ‘*Ca*. Uabimicrobiia’ strains has two possible explanations: the structure is absent, or it is present but undetected. For the sacculus, absence is more likely. TEM thin sections of ‘*Ca*. U. amorphum’ (Figure 3 b) show no wall-like layer, and the steric demands of engulfment in a diderm cell argue against a cell-spanning sacculus. For the division ring, the data does not favour either explanation. If the CDM II strains of *’Ca*. Uabimicrobiia’ divided without PG, however, predatory *Planctomycetota* would have three division mechanisms rather than two — CDM I, PG-dependent CDM II in HIEnr_7_s, and PG-independent CDM II in *’Ca*. Uabimicrobiia’. Since the CDM II strains of *’Ca*. Uabimicrobiia’ carry nearly the same complement of PG synthesis and remodelling genes as HIEnr_7_s, we consider a detection problem more likely than a third mechanism. Masking by the background fluorescence from feeding, a PG chemistry that excludes the D-amino acid probe, or remodelling too rapid to survive washing would each produce such a negative result. We have previously overturned the assumption that *Planctomycetota* lack a PG cell wall ^24^ and caution is warranted here as well. Separating the two explanations would require cryo-electron tomography. Chemical analysis of the envelope, the other decisive test, is precluded by the obligate co-culture with prey bacteria.

Shiratori and colleagues suggested that bacterial phagocytosis is analogous, not homologous, to eukaryotic phagocytosis and that a lokiactin homolog might act as a functional component in the process ^7^. The proposal was reasonable: actin-driven uptake is the core of eukaryotic phagocytosis, and the lokiactin gene appears to have reached ‘*Ca.* U. amorphum’ from Asgard archaea by horizontal gene transfer ^7^. Extending sequence- and structure-based searches to our previously unknown strains, however, we found the lokiactin homolog in only three of the nine BoP, ‘*Ca.* U. amorphum’ among them. A protein absent from six strains that engulf prey cannot be a general requirement of engulfment. Species-specific contributions in the strains carrying it, however, remain possible. Beyond the question of function, the presence of homologs of a gene of Asgard archaeal origin in two further BoP shows that such genes are not confined to ‘Ca. U. amorphum’ and can be found in BoP genomes more broadly. By contrast, the MCL proteins ^18^ fail the test for the opposite reason. They are conserved across all nine BoP, but equally present in the non-engulfing predator ‘*Ca.* N. helgolandensis’ and in other, non-predatory *Planctomycetota* ^15,19^. Their distribution seems to tie them to membrane remodelling in the phylum as a whole rather than to phagocytosis.

Neither search thus recovered a eukaryotic phagocytosis protein conserved specifically across the nine phagocytic strains. This absence admits two readings: an independent origin of bacterial phagocytosis, or an origin so remote that both sequence and structural signals have decayed beyond recovery. Our data do not distinguish between them; failure to detect remote homologues is a documented phenomenon ^25^, and the history of vesicle-coat protein identification counsels caution ^15,26,27^.

While the molecular machinery remains unknown, the FIB-SEM reconstructions settle a point at the cellular level that the first description had left open. The narrow ducts connecting some phagosome-like vacuoles to the exterior ^7^ are not a permanent feature: the mature vacuole is a closed compartment with no connection to the exterior, and the ducts most likely represent sections through cells captured during final stages of engulfment (Fig. 3c). The prey is therefore enclosed inside the predator in the strict sense. For this reason, we use the term bacterial phagocytosis rather than the earlier phagocytosis-like engulfment ^7^. The term describes the outcome, complete internalisation into a sealed compartment, not the mechanism, and implies no homology with the eukaryotic process. To our knowledge, this is the first demonstration that a prokaryote retains engulfed cells in a fully sealed intracellular compartment.

Finally, the isolates bear directly on questions concerning the processes whereby cells with complex architectures arise. Over the years, various models have been proposed for the origin of the eukaryotes, and these can broadly be sorted into mitochondria-first and mitochondria-late models. Mitochondria-first models rest on the premise that phagocytosis is energetically out of reach for a cell without mitochondria, so that the alphaproteobacterial endosymbiont must have been acquired by a non-phagocytic host ^9–11^. Shiratori et al. provided a first counterexample but, resting on a single enrichment culture, treated it strictly as a proof of principle, as did our own previous study ^8^. The strains described here move the point beyond proof of principle: whole-cell engulfment is a routine, aerobic lifestyle, performed across an entire planctomycetal class by cells without mitochondria and ending in a fully sealed compartment. One premise of mitochondria-first models is thus contradicted not by an isolated curiosity, but by a whole cultivated bacterial lineage.

These observations further raise the possibility that cells with complex intracellular network structures and phagocytic capabilities have evolved at least twice independently in both the bacterial and the archaeal/eukaryotic lineages. Various scenarios involving hosts from both lineages have been proposed for the origin of eukaryotes ^11,28–34^. Evaluating these scenarios will require cultivating further BoP as well as *Promethearchaeati* and making both lineages genetically accessible.

### Description of ‘*Candidatus* Nanovoracia’ classis nov

Na.no.vo.ra’ci.a. N.L. masc. n. *Nanovorax*, type genus of the class; N.L. neut. pl. n. *Nanovoracia*, the class of the genus *Nanovorax*. Cells are spherical. Predation via external digestion is observed. The type order is ‘*Ca*. Nanovoracales’. The class belongs to the phylum *Planctomycetota*.

### Description of ‘*Candidatus* Nanovoracales’ ord. nov

Na.no.vo.ra.ca’les. N.L. fem. pl. n. *Nanovoracales*, the order of the genus *Nanovorax.* Description as for the class. The type genus is ‘*Ca.* Nanovorax’.

### Description of ‘*Candidatus* Nanovoracaceae’ fam. nov

Na.no.vo.ra.ca’ce.ae. N.L. fem. pl. n. *Nanovoracaceae*, the family of the genus *Nanovorax.* Description as for the order. The type genus is ‘*Ca.* Nanovorax’.

### Description of ‘*Candidatus* Nanovorax’ gen. nov

Na.no.vo’rax. Gr. masc. n. *nanos*, dwarf; L. masc. adj. *vorax*, voracious; N.L. masc. n. *Nanovorax*, the small devourer of prey cells.

Aerobic, chemoheterotrophic, obligately predatory bacteria. Cells attach reversibly to prey bacteria and digest the prey externally without engulfment, leaving small rounded residual bodies. Division by binary-fission-like constriction via a transient peptidoglycan ring in the division plane at mid-cell. The genomic DNA G+C content is ∼52%. The type species is ‘*Ca.* Nanovorax helgolandensis’.

### Description of ‘*Candidatus* Nanovorax helgolandensis’ sp. nov

hel.go.lan.den’sis. N.L. masc. adj. *helgolandensis*, of Helgoland (the German name of Heligoland Island), the North Sea island from whose surrounding surface waters the organism was enriched.

Exhibits the properties of the genus. Cells are round with a diameter of 1-3 µm and grow at a temperature of 24 °C and a pH of 8.5; preys on *Alteromonas macleodii* in enrichment culture. The type strain is HIEnr_7_s. It was isolated from surface water in the harbour of Heligoland Island. The genome of the type strain has a size of 10.37 Mbp and the strain lacks extrachromosomal elements.

### Description of ‘*Candidatus* Ligurimicrobium’ gen. nov

Li.gu.ri.mi.cro’bi.um. N.L. masc. n. Ligur, a sea giant in the founding legend of the island of Heligoland; N.L. neut. n. microbium, a microbe; N.L. neut. n. *Ligurimicrobium*, Ligur’s microbe, referring to the size of the organism in comparison to its prey and to Heligoland, its site of isolation.

Aerobic and chemoheterotrophic cells with predatory lifestyle. Prey cells are internalized into fully sealed phagosome-like vacuoles (bacterial phagocytosis). Division by binary-fission-like constriction, distinguishing the genus from the other genera of the family, which divide by crawling apart on a surface until the emerging daughter cells are only connected by a thin, thread-like structure that eventually disrupts. Genomic DNA G+C content ∼35%. The type species is ‘*Ca.* Ligurimicrobium helgolandense’. The genus belongs to the family ‘*Ca.* Uabimicrobiaceae’, order ‘*Ca.* Uabimicrobiales,’ class ‘*Ca*. Uabimicrobiia’, phylum *Planctomycetota*.

### Description of ‘*Candidatus* Ligurimicrobium helgolandense’ sp. nov

hel.go.lan.den’se. N.L. neut. adj. *helgolandense*, of Helgoland, the German name for Heligoland Island.

Exhibits the properties of the genus. Cells are round with a diameter of ∼5 µm and grow at a temperature of 24 °C and a pH of 8.5; preys on *Alteromonas macleodii* in enrichment culture. The type strain is HlEnr_12. It was isolated from surface water in the harbour of Heligoland. The genome of the type strain has a size of 9.74 Mbp and the strain has three extrachromosomal elements.

### Description of ‘*Candidatus* Uabimicrobium palauense’ sp. nov

pa.lau.en’se. N.L. neut. adj. *palauense*, of Palau, the Pacific archipelago from whose waters the organism was enriched.

Cells are round with a diameter of ∼5 µm and grow at a temperature of 24 °C and a pH of 8.5; preys on *Alteromonas macleodii* in enrichment culture. The type strain is PEnr_61_6. It was isolated from surface water of the Wonder Lake in Palau. The genome of the type strain has a size of 7.96 Mbp and the strain lacks extrachromosomal elements.

## Materials and Methods

### Sampling, Isolation and Cultivation

Water samples were retrieved from various locations at the North Sea, the Baltic Sea, and the Pacific Ocean. For detailed sampling information see Supplementary Table 13. Samples were pre-enriched for 5 days in darkness at room temperature (20-21 °C) and subsequently distributed into petri-dishes for further enrichment. Cycloheximide (final concentration 50 mg/L) was added to all dish-enrichments, while LB Lennox (Sigma-Aldrich, Art. Nr. L3022, final concentration 0.001% (v/v)) was added to half of the enrichments originating from one sample. Enrichments were further incubated and growth of BoP was monitored using a cell culture microscope (VWR® VisiScope® IT415PH). Suspected BoP were transferred into new petri-dishes containing prey (*Alteromonas macleodii* DSM6062^T^) and sterile-filtered water from the sampling site, JCM Medium 1273 with artificial seawater from JCM Medium 1118 without LB, or TrM35 artificial sea water using a micro-pipette and were incubated in darkness at 24 °C. Subsequent cultures were maintained in the same manner. TrM35 artificial sea water (35 psu) consists of 40 g Tropic Marin® Classic sea salts (Tropic Marin, Art. Nr. 10112) dissolved in water (Milli-Q) to a final volume of 1 L. *A. macleodii* DSM 6062^T^ cultures were grown shaking overnight in Difco^TM^ Marine Broth 2216 (MB; BD, Art. Nr. 279110) at 28 °C. Cells were harvested by centrifugation for 5 min at 10,000 × g and washed three times in JCM Medium 1273 with artificial seawater from JCM Medium 1118 without LB. For feeding, the amount of *A. macleodii* cells added per 1 mL of medium equalled 2 / 4 µL of an *A. macleodii* cell suspension with OD_600_ = 15.0 for sterile filtered water from the sampling site or JCM Medium 1273 with artificial seawater from JCM Medium 1118 without LB, and TrM35 artificial sea water, respectively. The co-culture of ‘*Ca*. U. amorphum’ and *A. macleodii* was maintained as described previously ^8^. *E. coli* TOP10 and *E. coli* WM3064 were grown in LB Lennox supplemented with 0.3 mM 2,6-Diaminopimelic acid (DAP, Sigma-Aldrich, Art. Nr. 33240). Kanamycin was added for selection to a final concentration of 50 mg/L (*E. coli*) or 200 mg/L (*A. macleodii*). Conjugation was performed on TrM10PY-mateDAP medium (11.43 g Tropic Marin Sea Salts Classic, 10 g peptone, 5 g yeast extract per 1L water, pH adjusted to 7.5 with KOH) with 0.3 mM DAP. Solid media contained 15 g/L agar.

### Whole genome amplification and genome sequencing

Whole genome amplification (WGA) was performed using the REPLI-g Single Cell Kit from Qiagen as previously published ^8^. Briefly, cultures were grown in glass bottom dishes (Cellvis D35-10-1.5-N) until prey was almost completely depleted. Before lysis, cells were washed four times with culture medium of the respective strain (in case of sterile filtered environmental water, TrM35 or TrM10 (11.43 g Tropic Marin Sea Salts Classic per 1L water) were used instead followed by two washes with ddH2O. While lysis and DNA amplification was performed using double reaction volumes directly inside the glass bottom dish in most cases, few strains would not deplete the prey sufficiently. To improve the BoP:prey ratio, BoP cells were therefore scratched from a spot with little prey cells left. In this case WGA was performed in half reaction volumes using PCR tubes. Before heating the sample to 65 °C for 10 min, three additional incubations were performed at 4 °C, -20 °C, -80 °C (in this order) for 10 min each. DNA amplification was performed for 8 h. DNA purification and restriction of the obtained wgaDNA (whole genome amplified DNA) were performed following Oxford Nanopore’s Ligation sequencing gDNA – whole genome amplification protocol (version: WAL_9154_v112_revF_09Feb2022) with volumes during initial purification adjusted to the volumes of the WGA reaction as previously published ^8^. After binding the DNA to AMPure XP beads (Beckman Coulter), beads were pelleted on a magnet, washed with 70% (v/v) ethanol, and resuspended in nuclease-free water. The wgaDNA concentration was determined using the QubitTM 1x dsDNA BR kit (Thermo Scientific). Afterwards, wgaDNA was debranched as two separate samples using NEB T7 Endonuclease I with the incubation step at 37 °C being prolonged to 60 min. Subsequently, wgaDNA was bound to AMPure XP beads again, but using 90 µL AMPure XP bead suspension (Beckman Coulter) instead of 35 µL custom bead suspension and incubating on the Hula mixer for 20 min. After washing twice with 70% (v/v) ethanol and subsequent drying, the pellet was resuspended in 20 µL of nuclease-free water and the wgaDNA concentration was determined once again using a Qubit fluorometer.

Debranched wgaDNA was used to prepare a multiplex sequencing library for Oxford Nanopore long-read sequencing following the manufacturer’s protocol for the Native Barcoding Kit 24 (SQK-NBD114.24, Oxford Nanopore Technologies) with few modifications as reported before ^8^. Briefly, DNA repair was performed using NEBNext FFPE DNA Repair Mix and the NEBNext Ultra II End Repair / dA-tailing Module reagents (New England Biolabs) with incubation times at 65 °C and 20 °C each extended to 20 min. During the clean-up step with AMPure XP Beads (Beckman Coulter) the incubation times on the Hula mixer and after resuspension of the beads in nuclease-free water were extended to 10 min. Native barcodes were ligated to each wgaDNA sample using NEB Blunt/TA Ligase Mix (New England Biolabs) and the reaction was stopped after 30 min of incubation by adding EDTA. All samples were pooled and cleaned up with AMPure XP Beads (Beckman Coulter). Native sequencing adapters were ligated to the wgaDNA using the NEBNext Quick Ligation Module (New England Biolabs) for 30 min. Another purification step with AMPure XP Beads (Beckman Coulter) was performed with Long Fragment Buffer (Oxford Nanopore Technologies) instead of 80% (v/v) ethanol to enrich DNA fragments larger than 3 kb. The incubation time at 37 °C after resuspension of the beads in nuclease-free water was extended to 15 min. The final library was loaded on a primed R10.4.1 flow cell (FLO-MIN114, Oxford Nanopore Technologies) and sequencing was performed with a MinION Mk1B device (Oxford Nanopore Technologies). Raw nanopore sequencing data was basecalled using Dorado basecaller (versions and bascalling models are specified in the Supplementary Table 14 for each of the strains) with the additional flag “--min-qscore 10” for filtering reads with a Phred quality score ≥ 10 (Oxford Nanopore Technologies). Demultiplexing and adapter trimming were performed using Dorado demux. Additionally, wgaDNA was sequenced at Eurofins Genomics (Ebersberg, Germany) with an Illumina NovaSeq Sequencer in NovaSeq PE150 mode.

Nanopore long-read and Illumina short-read data were uploaded to the Galaxy web platform, and the server available under the public domain usegalaxy.eu was used for processing of the data using customized Galaxy workflows ^35^. The workflows included NanoPlot version 1.41.0 or version 1.43.0 ^36^ and FastQC version 0.74 ^37^ for quality control of the sequencing reads. Flye version 2.9.1 or version 2.9.5 was used for (meta-)genome assembly with the nanopore long reads (flag “--nano-hq”) ^38^. Assembly parameters were adapted using optional flags (“--meta”, “--scaffolds” or “--min-overlap”) where required to improve assembly quality (version numbers and the flags used for assembly are listed in Supplementary Table 14 for each of the strains). Medaka version 1.7.2 (Oxford Nanopore Technologies) with the model “r1041_e82_400bps_sup_g615” was used for polishing the raw assembly with the original long reads ^39^. The Illumina short reads were mapped to the medaka consensus sequences with BWA-MEM2 version 2.2.1 ^40,41^ and the mapping file was used for polishing with Pilon version 1.20.1 ^42^. The original long reads were mapped to the polished assembly with Minimap2 ^40,41,43^. The resulting mapping file was used to generate a tabular depth matrix file having mean and variance of the base coverage depth with MetaBAT2 version 2.15 or version 2.17 (version numbers are listed in Supplementary Table 14 for each of the strains) ^44^. Afterwards, polished contigs were binned using MetaBAT2 with the tabular depth matrix file (optional flag “--abdFile”). The resulting bins were annotated using Prokka version 1.14.6 ^45,46^. Based on the annotated 16S rRNA genes, bins were classified as BoPs (closest relative in the *Planctomycetota*), prey bacteria (*A. macleodii*) or contaminating bacterium of the enrichment (everything else). The contigs of each BoP bin were checked for circularity based on the Flye assembly report. Chromosomes were re-oriented to the start codon of the *dnaA* gene, encoding the replication initiator protein and plasmids were re-oriented to the start codon of the parA/soj gene, encoding an ATPase involved in chromosome/plasmid partitioning (if present on the plasmid). Re-annotation of the final assemblies was done using PGAP version 2026-06-18 build 8602 ^47^.

### Nucleotide sequence accession numbers

The 16S rRNA gene sequences were deposited in the GenBank database, while genome sequence information is available from NCBI under the accession numbers provided in Supplementary Table 13.

### Analyses of phylogenetic markers and tree construction

The full-length 16S rRNA and *rpoB* gene sequences were extracted from the prokka-annotated genomes, aligned with ClustalW^48^ and respective sequence similarity matrices were calculated with TaxonDC^49^ (for *rpoB* only the similarity of a ca. 1300 bp partial sequence of the gene was considered as previously described^50^). Average amino acid identities (AAI) and average nucleotide identities (ANI) were calculated using respective scripts of the enveomics v.1.10.0 collection (ANI was calculated as ANIb using BLAST) ^51,52^. The percentage of conserved proteins (POCP) was calculated as described^53^.

A database was generated from the protein sequences encoded by the predicted genes in the new *Planctomycetota* genomes. Orthologs to 114 proteins previously used as phylogenomic markers that resolved the diversification of early branching lineages in the *Planctomycetota* ^16^ were identified in BLASTp searches ^54^ against the database using proteins from ‘*Ca.* Uabimicrobium amorphum’ or ‘*Saltatorellus ferox*’ as the queries. If no hit was obtained, a tBLASTn search was used with the same query to search for genes that may not yet have been predicted in the new genomes. The expanded protein datasets were manually inspected using AliView to verify that the newly identified proteins were correct ^55^.

The species tree was computed following the procedure used in ^16^. In brief, each expanded protein dataset was aligned individually using MAFFT v7.471 (L-INS-I option)^56^. The resulting alignments were concatenated into a supermatrix using a custom Python script (concat_alignment.py), which also inserted gap characters for species missing from a given protein. The concatenated alignment was trimmed using BMGE v1.12 (-s FAST, -h 0:1 and -g 1) ^57^.

A guide tree was first computed under the LG+F+G model using IQ-TREE v1.6.12 ^58^. Branch supports were assessed using ultrafast bootstrap approximation (UFboot) with 1,000 bootstrap replicates ^59^ and SH-like approximative likelihood ratio test (SH-aLRT) ^60^, for which 1,000 replicates were used. This guide tree was then used to estimate per-site amino acid frequency profiles under the LG+C60+F+G mixture model via the Posterior Mean Site Frequency (PMSF) approach ^61^. Using the pre-computed site frequencies, 100 non-parametric bootstrap trees were generated independently under the LG+C60+F+G model. A consensus tree was constructed from the 100 bootstrap trees using IQ-TREE, and branch lengths were re-estimated on the consensus topology under the LG model. The tree was visualized using iTOL ^62^.

### Genome analyses

Biosynthetic gene clusters (BGCs) were predicted with antiSMASH v.8.0.4^63^ in relaxed mode and with all extra features activated. Carbohydrate-active enzymes (CAZymes) were analyzed with run_dbcan, the standalone tool of dbCAN3^64^. Gene origin was inferred with eggNOG-mapper v2.1.12 against eggNOG database version 5.0.2, Diamond version 2.0.11, MMseqs2 version 113e3212c137d026e297c7540e1fcd039f681 2b1 and compatible novel families database version 1.0.1 in protein search mode with default parameters (e-value 0.001, sensitive mode ^64^. For CAZyme domain/family annotation via HMMER an e-value of 1e-15 and a coverage cutoff value of 0.35 were used. The annotation with precomputed orthologous groups and phylogenies was performed with eggnog-mapper v. 2.1.12 ^65^.

To identify BoP’s components of the PG synthesis route, the divisome and the elongasome, the genomes were processed with anvi’o 8 ^66^. Open reading frames were predicted in anvi’o using Prodigal ^67^ and KOfams (KEGG Orthologs) ^68^ were identified using anvi-run-kegg-kofams. KOfam hits of each genome were exported from anvi’o and hits per KOfam to be analyzed were counted per genome. The data was plotted using RStudio ^69^ and ggplot2 ^70^.

Gene origin was inferred with eggNOG-mapper v2.1.12 against eggNOG database version 5.0.2, Diamond version 2.0.11, MMseqs2 version 113e3212c137d026e297c 7540e1fcd039f6812b1 and compatible novel families database version 1.0.1 in protein search mode with default parameters (e-value 0.001, sensitive mode) ^65^. Each query was assigned to the taxonomic level of its best-matching orthologous group. Assignments were visualized using RStudio^69,71^ with the packages readr^72^ (version 2.2.0), tidyr^73^ (version 1.3.1), dplyr^74^ (version 1.1.4), ggplot2^70^ and scales^75^. Genes were grouped by phylum and phyla by domain (Bacteria, Archaea, Eukarya and Viruses). Genes assigned generally to the domain Bacteria were retained as a separate category (“Bacteria (domain-level)”). Proportions were calculated per strain relative to all assigned genes including that category. Phyla not reaching 1% in any strain were pooled within their domain (“other”).

### Light microscopy

Light microscopy was performed using a Nikon Eclipse Ti2 inverted microscope equipped with a Nikon DS-Ri2 Camera for colored phase contrast images and a Hamamatsu ORCA-Flash4.0LT camera used for fluorescence images as well as black and white phase contrast images. A Nikon CFI Plan Apochromat Lambda 100X Ph3 oil immersion objective (numerical aperture 1.45) was employed for all light microscopy methods. Fluorescence microscopy was performed employing a SpectraX light source with pre/excitation-filters 395/25, 440/20, and 550/15 (CWL/BW). Feeding assays were imaged using a Semrock FF409/493/573/652-Di02 dicroic beamsplitter and a Semrock FF01-432/515/595/730 quad-band emission filter, HADA staining images were recorded using a DAPI-1160B (Semrock) filter set. Bleaching in time series was corrected using the Bleach correction ImageJ plugin in histogram matching mode ^76^. Afterwards, time series were stabilized using the ImageJ Image Stabilizer plugin ^77^. Brightness and contrast were adjusted to enhance visibility using FIJI (Version 2.14.0) ^78^. Cultures for non-fluorescence microscopy were grown as described above in glass bottom dishes (Ibidi 81158 or Cellvis D35-10-1.5-N). For preparation of fluorescent feeding assays, *A. macleodii* was prepared as described above, kanamycin was only added in the overnight culture. When working with both fluorescent strains, cells were mixed in a 1:1 (v:v) ratio and then added to a glass bottom dish containing JCM Medium 1273 with artificial seawater from JCM Medium 1118 without LB, TrM35 or TrM10 (11.43 g Tropic Marin Sea Salts Classic per 1L water) with 0.0075% (w/v) glucose. Afterwards, strains were added from a well-grown culture with no prey left at the bottom surface of the culture vessel. Cultures were imaged after incubation for 2-6 days. Cultures for staining with HADA (3-[[(7-Hydroxy-2-oxo-2*H*-1-benzopyran-3-yl)carbonyl]amino]-D-alanine hydrocholoride, Tocris Bioscience, Art. Nr. 6647/5) grown in glass bottom dishes were stained with 0.25 mM HADA (final concentration) for 5 h. Afterwards, cells were washed four times by flushing the glass bottom dish with JCM Medium 1273 with artificial seawater from JCM Medium 1118 without LB, TrM35 or TrM10.

### Electron microscopy

Cultures for scanning electron microscopy (SEM) were grown inside glass bottom dishes (Ibidi 81158), fixed for 30 min at room temperature with 1% glutaraldehyde (final concentration) and afterwards stored at 4 °C. The samples were transferred into 75 mM cacodylate buffer (pH 7.0) containing 2 mM MgCl_2_ and 300 mM NaCl and washed four times (10, 20, 30 and 90 min). Post-fixation was performed with 1% OsO_4_ for 1 hour. This was followed by one washing step with cacodylate buffer (10 min) and 3 washing steps with double-distilled water (10, 20 and 30 min). After dehydration in a graded acetone series, the samples were critical-point-dried, mounted on aluminium stubs and sputter-coated with platinum for 20 s. Scanning electron microscopy was carried out on a Zeiss Auriga 40 Crossbeam workstation (Zeiss, Oberkochen, Germany) at 1.5 kV acceleration voltage with signal detection using the SE-detector.

For the investigation via transmission electron microscopy (TEM), cells were scratched off petri-dishes and fixed with 1% glutaraldehyde in the above-mentioned cacodylate buffer. After centrifugation for 5 min at 5,000 x g, the pellet was resuspended in a small amount of buffer and mixed with 4% (w/v) low melt agarose and after solidification of the agarose covered again with fixative (1% glutaraldehyde in cacodylate buffer) for 1 h. The pellet was washed 4 times with cacodylate buffer (15, 30, 45 and 60 min) and post-fixed with 1% OsO_4_ for 1h. The pellet was then washed with cacodylate buffer overnight and three times with double-distilled water (15, 30 and 45 min). The cells were dehydrated in a graded acetone series including additional staining with 1% uranyl acetate for 20 min in the 20% step. The 100% acetone step was repeated three times (15, 60 and 60 min) before stepwise infiltration with Epon 812 epoxy resin and the final polymerization of the resin overnight at 63°C. With these samples, 70 nm thin sections were created and post-stained with 1% lead citrate for 2 min prior to electron microscopy. The samples were investigated on a JEOL F200 transmission electron microscopy (JEOL, Freising, Germany) at 200 kV. Images were recorded with a 6-megapixel EMSIS Tolara camera (EMSIS GmbH, Münster, Germany) and the Radius software package.

To get an impression of the 3D-architecture of the cells, focused ion beam scanning electron microscopy (FIB/SEM) was applied. Herein we used flat embedded cells on glass slides. For the latter, the fixation, washing and dehydration steps were identical to the TEM-preparation except that they were performed with cells adherent to glass slides in cuvettes. To infiltrate the cells with resin, droplets of rising concentrations of Epon 812 epoxy resin were put on the cells on the glass slide. After repeating the 100% resin step twice, the glass slides were put in a 50 mL Falcon Tube with a filter paper saturated with 100% acetone at the bottom and centrifuged for 2 min with 2,000 rpm to get rid of excessive resin and make the layer covering the cells thinner. The resin was then polymerized for 96 h at 63 °C, mounted on aluminium stubs and coated with 20 nm carbon.

FIB/SEM tomography was carried out on a Zeiss Auriga 40 crossbeam workstation. Gallium milling was carried out at a current of 50 pA with a layer thickness of 5 nm. By imaging after 3 or 4 milling rounds, we achieved z-resolutions of 15 or 20 nm. Imaging in between the milling steps was carried out at an acceleration voltage of 1.5 kV and signal detection with the energy selective backscatter (EsB) detector. For optimization of the images, we used Affinity Photo 2, followed by ImageJ for alignment and Amira^®^ (Thermo Fisher Scientific, Dreieich, Germany) for segmentation.

### Genetic manipulation of *Alteromonas macleodii* DSM6062

All cloning steps for construction of plasmids pCEW001-PgapAm-TurboRFP and pCEW001-PgapAm-GFPmut2 were carried out in *E. coli* TOP10. pCEW001 was constructed by amplification of a 3790 bp fragment from pBBR1MCS-2 ^79^ with primers 5’-TGTCTAGAGTGAAGCTTTCAGAATTCGTAATCGATTCAGGATCCTCACCTAGGAGGCGGCTACAGCCGATAGTCTGG-3’ and 5’-CTGTCTAGAACTGGTACCACTCTCGAGTCAGATATCCTGACGCGTACTCATATGACTGCTAGCTTCGAACCCCAGAGTCCCGCT CAG-3’, which was subsequently cut with XbaI (Thermo Scientific™, FD0685) and ligated using T4 Ligase (Thermo Scientific™, K1423). The native promoter of the glyceraldehyde-3-phosphate dehydrogenase gene of *A. macleodii* (P_gapAm_) was amplified using primers 5’-TCAGGATCCGGTGTGTCATTCCTGTTATTCGGTAAAAGAC-3’ and 5’-TCAGAATTCCCCCGATTATTTAAATAAAATGAAAATAAATTACGTAAAG-3’.

Subsequently, the amplified promoter was inserted into pCEW001 via the *Bam*HI and *Eco*RI restriction sites yielding pCEW001-P_gapAm_ (Enzymes: Thermo Scientific™ FD0055 and FD0274). Clones carrying plasmids with the correct inserts were confirmed by PCR with primers 5’-GCCGAAAAGTTGCAGCTGATTGCGCATG-3’ and 5’-GTGATATTGCTGAAGAGCTTGGCGGCGAATG-3’ and control digestion (EcoRV and BsaI; Thermo Scientific™ FD0303 and FD0294). Genes encoding TurboRFP and GFPmut2 were amplified with primers 5’-TCAGAATTCAAGGAGATCATTCATGGGAAGTGAATTGATTAAAGAAAATATGCATATGAAATTATATATG-3’ and 5’-CTGAAGCTTTTATCTATGCCCTAATTTACTAGGTAAATCACAATATTTTGCTAC-3’ (TurboRFP) and 5’-TCAGAATTCAAGGAGATCATTCATGAGTAAAGGAGAAGAACTTTTCACTGGAGTTGTCCCAATTC-3’ and 5’-CTGAAGCTTTTATTTGTATAGTTCATCCATGCCATGTGTAATCCCAGC-3’ (GFPmut2), respectively. Fragments were inserted via the *Eco*RI and *Hind*III restriction sites to yield pCEW001-P_gapAm_-TurboRFP and pCEW001-P_gapAm_-GFPmut2 (Enzymes: Thermo Scientific™ FD0274 and FD0505). Plasmids carrying the correct inserts were confirmed by PCR with the same primers as for plasmid pCEW001-P_gapAm_ and control digestion (pCEW001-P_gapAm_-TurboRFP: XbaI, Thermo Scientific™ FD0685; pCEW001-P_gapAm_-GFPmut2, NcoI, Thermo Scientific™ FD0574). Plasmids were transferred via heat shock into *E. coli* WM3064 for conjugation into *A. macleodii*. The conjugation protocol was adapted from ^80^ and ^81^. Overnight cultures were grown of both donor and recipient. 2 mL recipient and 1 mL donor culture were harvested by centrifugation (5000 × g, 2 min) and washed twice in TrM10PY-mateDAP. Recipient and donor were mixed by resuspension in 100 µL TrM10YP-mateDAP and spotted onto TrM10YP-mateDAP plates. After incubation over night at 28 °C, the emerged lawn was collected into 1 mL MB. Cells were centrifuged at 5,000 × g for 2 min and subsequently wash twice in MB before plating on MB plates containing kanamycin. Plates were incubated overnight at 28 °C and colonies carrying the plasmid were confirmed via PCR targeting the fluorescence gene and P_gapAm_.

### Comparative genomic approaches

Based on Supplementary Dataset 7 of ^82^, a curated List of 192 eukaryotic phagocytosis genes was obtained by filtering the clade origin for ‘Eukaryota’ and the origin in phagocytosis for ‘Phagotrophy’. Afterwards, functions ‘Apoptosis/Autophagy’, ‘Chaperone’, ‘Hydrolases’, ‘Immunity/Inflammation’, ‘Membrane Receptor’, ‘Metabolism’, ‘Metabolism (lipid)’, ‘Metabolism (Mitochondrial)’, ‘Nucleotide associated protein’, ‘Proteasome’, ‘Protein Biosynthesis’, ‘Protein metabolism’, ‘RNA binding’, ‘Signaling’, ‘small GTPase’, ‘Transporter (Mitochondrial)’, ‘Transporter (Transmembrane)’ were omitted leaving Proteins associated with functions ‘Cellular trafficking’, ‘Cytoskeleton’, ‘Membrane Structure’, and ‘Unknown’ (Supplementary Table 9).

To identify gene families potentially involved in phagocytosis and the different cell division modes, we gathered a dataset of 104 *Planctomycetota* MAGs and genomes. This includes the genomes reported in this study, as well as genomes of representatives of non-phagocyting *Planctomycetota* as outgroups (four genome representatives per *Planctomycetota* GTDB classes). The orthologous gene groups were clustered using OrthoFinder ^83,84^ pipeline (version 2.5.5), specifying the use of Diamond (v2.0.15.153) ^85^ for protein similarity search, multiple sequence alignment for tree inferences and splitting paralogous clades bellow root (-S diamond -M msa -y). The gene families that were shared between phagocyting representatives and were absent from non-phagocyting representatives were subsequently functionally annotated using both sequence and structural based annotations. The structure of these proteins was predicted using AlphaFold3 ^86^. Then, the best structural predictions were used for structural based functional annotation using Foldseek (version 10.941cd33) ^87^ online against multiple databases (PDB, UniProt50, SwissProt, Proteome, BFVDv2 and MGnify). Additionally, eggNOG mapper (version 2.1.12) was used for sequence based functional annotation against eggNOG DB (version: 5.0.2) ^65^.

### Identification of membrane coat-like proteins

The predicted proteomes of the nine previously unknown strains were compiled into a local database and membrane coat-like (MCL) proteins were identified using two complementary approaches. First, bacterial MCL proteins were used as BLASTp queries (BLAST+ v2.12.0; ^88^): the MCL protein of ‘*Ca.* Uabimicrobium amorphum’ (UniProt A0A5S9IMS2), a homologue of *Gemmata obscuriglobus* MCL19, and the 19 MCL proteins described for *G. obscuriglobus* ^15^. Hits with an E-value < 10⁻⁴ were considered MCL candidates. Second, to detect more distant eukaryotic homologs, profile hidden Markov models (HMMs) were built from six membrane coat (MC) proteins of *Saccharomyces cerevisiae*: clathrin heavy chain (UniProt P22137), clathrin light chain (P17891), COPI α-subunit (P53622), the nucleoporins Nup85 (P46673) and Nup133 (P36161), and the COPII outer-coat protein Sec31 (P38968). For each query, jackhmmer (HMMER v3.1b2; ^89^) was run for 3 iterations against a local database of eukaryotic proteomes (PMID: 40033103). The resulting HMMs were used as queries against the strain proteomes with hmmsearch (E-value < 10^-4^). Hits from both approaches were merged and duplicates removed.

### Structure prediction of MCL candidates

Structures of all candidate proteins were predicted using ColabFold v1.5.5 ^90^ (MSA mode, number of models = 3, recycles 1 or 3) or AlphaFold 3 server ^86^. A candidate was accepted as an MCL protein only if its predicted model showed the architecture typical of MC proteins: β-propeller domains combined with an α-solenoid ^15,18^, assessed by visual inspection.

### Remote homology search for eukaryotic phagocytosis proteins

To test whether components of the eukaryotic phagocytic machinery are encoded in the BoP, we used 191 *Mus musculus* phagosome proteins ^82^ as queries (full list in Supplementary Table 9). For each query, a profile HMM was generated by three iterations of jackhmmer against the local eukaryotic database described above. The 191 HMMs were then searched against the proteomes of the nine strains with hmmsearch (E-value < 10^-4^). A query was scored as present in a strain if it contained at least one hit below the threshold.

## Author contributions

C.J. conceived and supervised the study. C.E.W. and V.H. collected environmental samples. C.E.W. isolated and cultivated the strains, performed prey-uptake experiments and HADA peptidoglycan labelling. C.E.W. genetically modified prey bacteria with help from T.H. and N.K.. N.K., C.E.W. and T.H. performed the eggNOG-based analysis of horizontal gene transfer. T.H. and C.E.W. performed light microscopy. A.K. and J.M.M. performed electron microscopy and FIB-SEM 3D reconstruction. J.H. performed genome sequencing and assembly. N.K. performed genome annotation and upload. S.G.E.A., B.S.B., A.J.G. and N.K.performed the phylogenomic analyses. M.T. and K.K. retrieved the groundwater metagenome-assembled genomes and computed the phylogeny linking them to the isolates. D.D., M.L.F. and L.F.M.-M. analyzed membrane-coat-like proteins and conducted the targeted search for eukaryotic phagocytosis proteins, sequence-based search and confirmed by structure predictions. A.J.P., F.B. and A.R.S. performed the untargeted structure-based homology search. C.E.W., A.J.P. and M.L.F. analyzed cell-division genes. T.H. contributed to protein structure predictions. C.J. wrote the manuscript with help from C.E.W. and from all authors. All authors read and approved the final manuscript.

## Acknowledgment

We thank René Maskos, Nicole Wohlfarth and Hanan Alderzy for skillful technical assistance. We further thank the Biological Institute Helgoland, Alfred Wegener Institute, and Helmholtz Centre for Polar and Marine Research for supporting our sampling campaign. Additionally, we thank the groups of Prof. Dr. Kai Papenfort (Microbial Gene Regulation, Friedrich Schiller University Jena) and Dr. Kathrin Fröhlich (Bacterial RNA Biology, Friedrich Schiller University Jena) for providing the plasmid harboring the TurboRFP encoding gene. We thank Dr. David Moyano (CABIMER, Sevilla, SP) for structural predictions in the positive screen. We further thank Ketil Krabbe, Kevin Bisdorf and Josefine Bach for technical assistance with groundwater enrichments.

## References

1 Perez, J., Moraleda-Munoz, A., Marcos-Torres, F. J. & Munoz-Dorado, J. Bacterial predation: 75 years and counting! Environ Microbiol 18, 766–779 (2016). 10.1111/1462-2920.13171

2 Guerrero, R. et al. Predatory prokaryotes: predation and primary consumption evolved in bacteria. Proceedings of the National Academy of Sciences of the United States of America 83, 2138–2142 (1986). 10.1073/pnas.83.7.2138

3 Moreira, D., Zivanovic, Y., Lopez-Archilla, A. I., Iniesto, M. & Lopez-Garcia, P. Reductive evolution and unique predatory mode in the CPR bacterium *Vampirococcus lugosii*. Nat Commun 12, 2454 (2021). 10.1038/s41467-021-22762-4

4 Yakimov, M. M. et al. Cultivation of a vampire: ‘*Candidatus* Absconditicoccus praedator’. Environ Microbiol 24, 30–49 (2022). 10.1111/1462-2920.15823

5 Stolp, H. & Starr, M. P. *Bdellovibrio bacteriovorus* gen. et sp. n., a predatory, ectoparasitic, and bacteriolytic microorganism. Antonie van Leeuwenhoek 29, 217–248 (1963). 10.1007/BF02046064

6 Berleman, J. E. & Kirby, J. R. Deciphering the hunting strategy of a bacterial wolfpack. FEMS Microbiol Rev 33, 942–957 (2009). 10.1111/j.1574-6976.2009.00185.x

7 Shiratori, T., Suzuki, S., Kakizawa, Y. & Ishida, K. I. Phagocytosis-like cell engulfment by a planctomycete bacterium. Nat. Commun. 10, 5529 (2019). 10.1038/s41467-019-13499-2

8 Wurzbacher, C. E. et al. “*Candidatus* Uabimicrobium helgolandensis”-a planctomycetal bacterium with phagocytosis-like prey cell engulfment, surface-dependent motility, and cell division. mBio, e0204424 (2024). 10.1128/mbio.02044-24

9 Lane, N. & Martin, W. The energetics of genome complexity. Nature 467, 929–934 (2010). 10.1038/nature09486

10 Martin, W. F., Garg, S. & Zimorski, V. Endosymbiotic theories for eukaryote origin. Philos Trans R Soc Lond B Biol Sci 370, 20140330 (2015). 10.1098/rstb.2014.0330

11 Mills, D. B. The origin of phagocytosis in Earth history. Interface Focus 10, 20200019 (2020). 10.1098/rsfs.2020.0019

12 Wiegand, S., Jogler, M. & Jogler, C. On the maverick *Planctomycetes*. FEMS Microbiol Rev 42, 739–760 (2018). 10.1093/femsre/fuy029

13 Rivas-Marin, E. & Devos, D. P. The Paradigms They Are a-Changin’: past, present and future of PVC bacteria research. Antonie Van Leeuwenhoek 111, 785–799 (2018). 10.1007/s10482-017-0962-z

14 Santarella-Mellwig, R. et al. The compartmentalized bacteria of the *Planctomycetes*-*Verrucomicrobia*-*Chlamydiae* superphylum have membrane coat-like proteins. PLoS Biol 8, e1000281 (2010). 10.1371/journal.pbio.1000281

15 Ferrelli, M. L. et al. Prokaryotic membrane coat - like proteins: An update. Journal of structural biology 215, 107987 (2023). 10.1016/j.jsb.2023.107987

16 Odelgard, A., Hagglund, E., Guy, L. & Andersson, S. G. E. Phylogeny and expansion of serine/threonine kinases in phagocytotic bacteria in the phylum *Planctomycetota*. Genome biology and evolution 16 (2024). 10.1093/gbe/evae068

17 Viklund, J., Ettema, T. J. & Andersson, S. G. Independent genome reduction and phylogenetic reclassification of the oceanic SAR11 clade. Molecular biology and evolution 29, 599–615 (2012). 10.1093/molbev/msr203

18 Devos, D. et al. Components of coated vesicles and nuclear pore complexes share a common molecular architecture. PLoS Biol 2, e380 (2004). 10.1371/journal.pbio.0020380

19 Lonhienne, T. G. et al. Endocytosis-like protein uptake in the bacterium *Gemmata obscuriglobus*. Proceedings of the National Academy of Sciences of the United States of America 107, 12883–12888 (2010). 10.1073/pnas.1001085107

20 Santarella-Mellwig, R. et al. The compartmentalized bacteria of the *Planctomycetes-Verrucomicrobia-Chlamydiae* superphylum have membrane coat-like proteins. PLoS Biol 8 (2010). 10.1371/journal.pbio.1000281

21 Liechti, G. W. et al. A new metabolic cell-wall labelling method reveals peptidoglycan in *Chlamydia trachomatis*. Nature 506, 507–510 (2014). 10.1038/nature12892

22 Liechti, G. et al. Pathogenic *Chlamydia* lack a classical sacculus but synthesize a narrow, mid-cell peptidoglycan ring, regulated by MreB, for cell division. PLoS pathogens 12, e1005590 (2016). 10.1371/journal.ppat.1005590

23 Pilhofer, M. et al. Discovery of chlamydial peptidoglycan reveals bacteria with murein sacculi but without FtsZ. Nat Commun 4, 2856 (2013). 10.1038/ncomms3856

24 Jeske, O., et al. *Planctomycetes* do possess a peptidoglycan cell wall. Nat Commun 6, 7116 (2015). 10.1038/ncomms8116

25 Weisman, C. M., Murray, A. W. & Eddy, S. R. Many, but not all, lineage-specific genes can be explained by homology detection failure. PLoS Biol 18, e3000862 (2020). 10.1371/journal.pbio.3000862

26 Akil, C. & Robinson, R. C. Genomes of *Asgard* archaea encode profilins that regulate actin. Nature 562, 439–443 (2018). 10.1038/s41586-018-0548-6

27 Boedeker, C. et al. Determining the bacterial cell biology of *Planctomycetes*. Nat Commun 8, 14853 (2017). 10.1038/ncomms14853

28 Devos, D. P. & Reynaud, E. G. Evolution. Intermediate steps. Science 330, 1187–1188 (2010). 10.1126/science.1196720

29 Fuerst, J. A. & Sagulenko, E. Beyond the bacterium: *Planctomycetes* challenge our concepts of microbial structure and function. Nat Rev Microbiol 9, 403–413 (2011). 10.1038/nrmicro2578

30 Devos, D. P. Reconciling *Asgardarchaeota* phylogenetic proximity to eukaryotes and *Planctomycetes* cellular features in the evolution of life. Molecular biology and evolution 38, 3531–3542 (2021). 10.1093/molbev/msab186

31 Eme, L. et al. Inference and reconstruction of the heimdallarchaeial ancestry of eukaryotes. Nature 618, 992–999 (2023). 10.1038/s41586-023-06186-2

32 Imachi, H. et al. Isolation of an archaeon at the prokaryote-eukaryote interface. Nature 577, 519–525 (2020). 10.1038/s41586-019-1916-6

33 Lopez-Garcia, P. & Moreira, D. Eukaryogenesis, a syntrophy affair. Nat Microbiol 4, 1068–1070 (2019). 10.1038/s41564-019-0495-5

34 Vosseberg, J. et al. The emerging view on the origin and early evolution of eukaryotic cells. Nature 633, 295–305 (2024). 10.1038/s41586-024-07677-6

35 Community, T. G. Galaxy for accessible, reproducible, and collaborative data analyses: 2026 update. Nucleic acids research 54, W105–W116 (2026). 10.1093/nar/gkag469

36 De Coster, W., D’Hert, S., Schultz, D. T., Cruts, M. & Van Broeckhoven, C. NanoPack: visualizing and processing long-read sequencing data. Bioinformatics 34, 2666–2669 (2018). 10.1093/bioinformatics/bty149

37 Andrews, S. FastQC: a quality control tool for high throughput sequence data. (Babraham Bioinformatics, 2010).

38 Lin, Y. et al. Assembly of long error-prone reads using de Bruijn graphs. Proceedings of the National Academy of Sciences 113, E8396–E8405 (2016). doi:10.1073/pnas.1604560113

39 Ltd., O. N. T. medaka: Neural-network consensus polishing and variant calling for Oxford Nanopore sequencing data. (2020).

40 Li, H. & Durbin, R. Fast and accurate short read alignment with Burrows-Wheeler transform. Bioinformatics 25, 1754–1760 (2009). 10.1093/bioinformatics/btp324

41 Li, H. & Durbin, R. Fast and accurate long-read alignment with Burrows–Wheeler transform. Bioinformatics 26, 589–595 (2010). 10.1093/bioinformatics/btp698

42 Walker, B. J. et al. Pilon: an integrated tool for comprehensive microbial variant detection and genome assembly improvement. PloS one 9, e112963 (2014). 10.1371/journal.pone.0112963

43 Li, H. Minimap2: pairwise alignment for nucleotide sequences. Bioinformatics 34, 3094–3100 (2018). 10.1093/bioinformatics/bty191

44 Kang, D. D. et al. MetaBAT 2: an adaptive binning algorithm for robust and efficient genome reconstruction from metagenome assemblies. PeerJ 7, e7359 (2019). 10.7717/peerj.7359

45 Cuccuru, G. et al. Orione, a web-based framework for NGS analysis in microbiology. Bioinformatics 30, 1928–1929 (2014). 10.1093/bioinformatics/btu135

46 Seemann, T. Prokka: rapid prokaryotic genome annotation. Bioinformatics 30, 2068–2069 (2014). 10.1093/bioinformatics/btu153

47 Tatusova, T. et al. NCBI prokaryotic genome annotation pipeline. Nucleic acids research 44, 6614–6624 (2016). 10.1093/nar/gkw569

48 Thompson, J. D., Gibson, T. J. & Higgins, D. G. Multiple sequence alignment using ClustalW and ClustalX. Current Protocols in Bioinformatics 00, 2-3 1–2 3 22 (2003).

49 Tarlachkov, S. & Starodumova, I. TaxonDC: Calculating the similarity value of the 16S rRNA gene sequences of prokaryotes or ITS regions of fungi. Journal of Bioinformatics and Genomics 3, 1–4 (2017).

50 Bondoso, J., Harder, J. & Lage, O. M. *rpoB* gene as a novel molecular marker to infer phylogeny in *Planctomycetales*. Antonie Van Leeuwenhoek 104, 477–488 (2013).

51 Rodriguez-R, L. M. & Konstantinidis, K. T. The enveomics collection: a toolbox for specialized analyses of microbial genomes and metagenomes; 2167-9843. PeerJ Preprints 4, e1900v1901 (2016).

52 Goris, J. et al. DNA-DNA hybridization values and their relationship to whole-genome sequence similarities. Int J Syst Evol Microbiol 57, 81–91 (2007). 10.1099/ijs.0.64483-0

53 Qin, Q. L., et al. A proposed genus boundary for the prokaryotes based on genomic insights. Journal of Bacteriology 196, 2210–2215 (2014).

54 Altschul, S. F. et al. Gapped BLAST and PSI-BLAST: a new generation of protein database search programs. Nucleic acids research 25, 3389–3402 (1997).

55. Larsson, A. AliView: a fast and lightweight alignment viewer and editor for large datasets. Bioinformatics 30, 3276-3278 (2014). 10.1093/bioinformatics/btu531

56 Katoh, K. & Standley, D. M. MAFFT multiple sequence alignment software version 7: improvements in performance and usability. Molecular biology and evolution 30, 772–780 (2013). 10.1093/molbev/mst010

57 Criscuolo, A. & Gribaldo, S. BMGE (Block Mapping and Gathering with Entropy): a new software for selection of phylogenetic informative regions from multiple sequence alignments. BMC evolutionary biology 10, 210 (2010).

58 Nguyen, L.-T., Schmidt, H. A., Von Haeseler, A. & Minh, B. Q. IQ-TREE: a fast and effective stochastic algorithm for estimating maximum-likelihood phylogenies. Molecular biology and evolution 32, 268–274 (2015).

59 Hoang, D. T., Chernomor, O., von Haeseler, A., Minh, B. Q. C Vinh, L. S. UFBoot2: Improving the Ultrafast Bootstrap Approximation. Molecular biology and evolution 35, 518–522 (2018). 10.1093/molbev/msx281

60 Guindon, S. et al. New Algorithms and Methods to Estimate Maximum-Likelihood Phylogenies: Assessing the Performance of PhyML 3.0. Systematic Biology 59, 307–321 (2010). 10.1093/sysbio/syq010

61 Wang, H.-C., Minh, B. Q., Susko, E. & Roger, A. J. Modeling site heterogeneity with posterior mean site frequency profiles accelerates accurate phylogenomic estimation. Systematic biology 67, 216–235 (2018).

62 Letunic, I. & Bork, P. Interactive Tree of Life (iTOL) v6: recent updates to the phylogenetic tree display and annotation tool. Nucleic acids research 52, W78–W82 (2024). 10.1093/nar/gkae268

63 Blin, K. et al. antiSMASH 8.0: extended gene cluster detection capabilities and analyses of chemistry, enzymology, and regulation. Nucleic Acids Research 53, W32–W38 (2025). 10.1093/nar/gkaf334

64 Zheng, J. et al. dbCAN3: automated carbohydrate-active enzyme and substrate annotation. Nucleic Acids Research 51, W115–W121 (2023). 10.1093/nar/gkad328

65 Cantalapiedra, C. P., Hernández-Plaza, A., Letunic, I., Bork, P. & Huerta-Cepas, J. eggNOG-mapper v2: Functional Annotation, Orthology Assignments, and Domain Prediction at the Metagenomic Scale. Molecular biology and evolution 38, 5825–5829 (2021). 10.1093/molbev/msab293

66 Eren, A. M. et al. Community-led, integrated, reproducible multi-omics with anvi’o. Nature Microbiology 6, 3–6 (2021). 10.1038/s41564-020-00834-3

67 Hyatt, D. et al. Prodigal: prokaryotic gene recognition and translation initiation site identification. BMC Bioinformatics 11, 119 (2010). 10.1186/1471-2105-11-119

68 Kanehisa, M. & Goto, S. KEGG: Kyoto Encyclopedia of Genes and Genomes. Nucleic Acids Research 28, 27–30 (2000). 10.1093/nar/28.1.27

69 Posit team, RStudio: Integrated Development Environment for R. Posit Software, PBC, Boston, MA. URL http://www.posit.co/. 2024).

70 Wickham, H. ggplot2: Elegant Graphics for Data Analysis, Springer-Verlag New York, ISBN 978-3-319-24277-4, https://ggplot2.tidyverse.org. (2016).

71 R: A Language and Environment for Statistical Computing. (R Foundation for Statistical Computing, Vienna, Austria, 2024).

72 readr: Read Rectangular Text Data v. R package version 2.2.0 (Wickham H, Hester J, Bryan J, 2026).

73 tidyr: Tidy Messy Data. v. R package version 1.3.1 (Wickham H, Vaughan D, Girlich M, 2024).

74 dplyr: A Grammar of Data Manipulation. v. R package version 1.1.4 (Wickham H, François R, Henry L, Müller K, Vaughan D, 2023).

75 scales: Scale Functions for Visualization v. R package version 1.4.0 (Wickham H, Pedersen T, Seidel D, 2025).

76 Miura, K. Bleach correction ImageJ plugin for compensating the photobleaching of time-lapse sequences. F1000Research 9 (2020). 10.12688/f1000research.27171.1

77. The image stabilizer plugin for ImageJ (2008).

78 Schindelin, J. et al. Fiji: an open-source platform for biological-image analysis. Nature methods 9, 676–682 (2012). 10.1038/nmeth.2019

79 Kovach, M. E., Phillips, R. W., Elzer, P. H., Roop, R. M., 2nd & Peterson, K. M. pBBR1MCS: a broad-host-range cloning vector. BioTechniques 16, 800–802 (1994).

80 Wang, Y. et al. Development of a transformation system for *Chlamydia trachomatis*: restoration of glycogen biosynthesis by acquisition of a plasmid shuttle vector. PLoS pathogens 7, e1002258 (2011). 10.1371/journal.ppat.1002258

81 Jung, A. et al. Molecular toolbox for genetic manipulation of the stalked budding bacterium *Hyphomonas neptunium*. Applied and environmental microbiology 81, 736–744 (2015). 10.1128/aem.03104-14

82 Boulais, J. et al. Molecular characterization of the evolution of phagosomes. Molecular systems biology 6, MSB201080 (2010). 10.1038/msb.2010.80

83 Emms, D. M. & Kelly, S. OrthoFinder: phylogenetic orthology inference for comparative genomics. Genome biology 20, 238 (2019). 10.1186/s13059-019-1832-y

84 Emms, D. M. & Kelly, S. OrthoFinder: solving fundamental biases in whole genome comparisons dramatically improves orthogroup inference accuracy. Genome biology 16, 157 (2015). 10.1186/s13059-015-0721-2

85 Buchfink, B., Reuter, K. & Drost, H.-G. Sensitive protein alignments at tree-of-life scale using DIAMOND. Nature methods 18, 366–368 (2021). 10.1038/s41592-021-01101-x

86 Abramson, J. et al. Accurate structure prediction of biomolecular interactions with AlphaFold 3. Nature 630, 493–500 (2024). 10.1038/s41586-024-07487-w

87 van Kempen, M. et al. Fast and accurate protein structure search with Foldseek. Nature biotechnology 42, 243–246 (2024). 10.1038/s41587-023-01773-0

88 Altschul, S. F., Gish, W., Miller, W., Myers, E. W. & Lipman, D. J. Basic local alignment search tool. Journal of molecular biology 215, 403–410 (1990). 10.1016/S0022-2836(05)80360-2

89 Eddy, S. R. Accelerated Profile HMM Searches. PLoS computational biology 7, e1002195 (2011). 10.1371/journal.pcbi.1002195

90 Mirdita, M. et al. ColabFold: making protein folding accessible to all. Nature methods 19, 679–682 (2022). 10.1038/s41592-022-01488-1

