## Supplementary Information for "Phagocytosis without mitochondria is a lineage trait of planctomycetal bacteria of prey"

### 1. Supplementary Notes

#### Bacteria of prey MAGs from groundwater

The occurrence of predatory *Planctomycetota* does not appear to be limited to marine systems. We were also able to detect MAGs from related organisms in enrichment cultures obtained from groundwater of carbonate rock aquifers, indicating that predatory *Planctomycetota* might be widespread in aquatic habitats. Phylogenomic analysis placed the groundwater MAGs into a group of *Planctomycetota* distinct from '*Ca.* Uabimicrobiia', and in closer relationship to '*Ca.* Nanovorax helgolandensis' HlEnr_7_s within class c__J058. Likewise, the DNA G+C contents of these MAGs are in the range of 56 to 70%, distinctly higher than in '*Ca.* Uabimicrobiia', while in the range of strain HlEnr_7_s. Although evidence for feeding on other bacteria has yet to be established due to the lack of cultures, if sharing the non-phagocytotic but predatory lifestyle of HlEnr_7_s, these organisms might be adapted to feed on cells from microcolonies on rock surfaces [1,2]. These might provide a more consistent source of prey than planktonic bacteria, given the low cell number of $10^4$ to $10^5$ $mL^{-1}$ present in the groundwater [3,4]. Furthermore, a surface-attached lifestyle might also be supported by the observed motility of predatory *Planctomycetota* and the attachment-based feeding mode of HlEnr_7_s.

### 2. Supplementary Methods

**Predatory *Planctomycetota* MAGs from groundwater**

Setup and incubation of enrichments

Groundwater was sampled in March 2021 from a well H41 (51.1150842 N, 10.4479713 E, 48 m sampling depth) of the Hainich Critical Zone Exploratory (CZE) in western Thuringia, Germany [5]. Water from this well features dysoxic conditions (4.83 mg/L $O_2$, 0.08 ± 0.08 mg/L $NH_4^+$, 8.5 ± 4.9 mg/L $NO_3^-$, 91.2 ± 28.9 mg/L $SO_4^{2-}$, pH 7.2) [6]. Water was pumped using a submersible motor pump (MP1, Grundfos, Denmark) until physicochemical parameters stabilized and subsequently collected in a 10 L high-density polyethylene carboy. After transfer to the laboratory and storage at 15 °C, triplicate incubations were set up the following week in autoclaved 1 L glass bottles with 750 mL groundwater and 2 mM methylamine (final concentration). Bottles were closed with air-tight silicon stoppers and incubated at 15 °C under mild shaking (50 rpm). Every nine to twelve months, the enrichments were passaged by adding 50 mL of enrichment to 700 mL of freshly sampled groundwater and supplying with 2 mM methylamine (final concentration).

Sampling and DNA extraction for metagenomics sequencing

Samples for metagenomic sequencing were taken from the fourth passage of the enrichments started on 4$^{th}$ October 2022. Samples were taken on 4$^{th}$ January 2023 from all replicates and on 14$^{th}$ December 2022 from enrichment replicate B. For each sample, 50 mL of the corresponding enrichment were filtered onto a 0.1 µm pore size PES filter (Merck, Darmstadt, Germany). DNA extraction was subsequently conducted using the

DNeasy PowerWater Kit (QIAGEN, Hilden, Germany) following the manufacturer's instructions.

Metagenomics sequencing

The quality of extracted DNA was checked on a 4200 TapeStation System (Agilent, Santa Clara, CA, USA) using the Genomic DNA ScreenTape (Agilent). Paired-end libraries were generated with an NEBNext Ultra II FS DNA preparation kit (New England Biolabs, Ipswich, MA, USA), and sequenced (2×150 bp) on an Illumina NovaSeq 6000 system (Illumina, San Diego, CA, USA) at the Fritz Lipmann Institute (Jena, Germany). Sequencing output ranged from 20.6 to 25.7 Gbp per sample, with an average of 23.0 Gbp.

Generation of metagenome-assembled genomes

The reads obtained from metagenomic sequencing were subjected to quality control using BBDuk (v39.01) [7]. Quality-controlled reads were subsequently assembled using metaSPAdes (v3.15.2) [8]. Genome binning was carried out using five binning algorithms: CONCOCT (v.1.0.0) [9], metabat2 (v2.12.1) [10] and MaxBin2 (v.2.2.6) [11] were run via metaWRAP (v1.3.2) [12] using contigs of 1000 bp and above. Both the 40 and 107 marker gene sets were utilized in MaxBin2. Furthermore, BinSanity (v0.2.7) [13] and Abawaca (v1.0.7) [14] were used to generate bins using contigs of 3000 bp and above. For Abawaca, 4-mer frequencies calculated using contigs with a minimal length of 5000 bp and 10000 bp were used. The generated bins were subsequently refined using metaWRAP, with thresholds of ≥50% completeness and ≤10% contamination. For dereplication of bins across samples, dRep (v3.4.0) [15] was used with a cutoff of 99% average nucleotide identity (ANI) to remove strain-level redundancy. Subsequently, quality of the final MAGs was assessed using checkM (v.1.2.2) [16], based on the presence of single copy marker genes. MAGs with at least 50% genome completeness and a maximum of 10% redundancy/contamination were then selected for analysis.

<u>Taxonomic characterization of MAGs</u>

Taxonomic classification of MAGs was conducted via GTDB-Tk (v2.3.0) [17], using the genome taxonomy database (release 214) as reference database. MAGs were screened via 16S rRNA gene sequences of known predatory *Planctomycetota* and three such MAGs were found. Relative abundances of MAGs in individual samples were calculated based on genome coverages determined using coverM (v.0.6.1) [18], following normalization based on total reads per sample. Phylogenomic comparison of the obtained groundwater MAGs with the genomes of the isolated bacteria of prey and further related *Planctomycetota* obtained from NCBI were conducted following the anvi'o workflow (v9) [19], based on a set of 71 bacterial single copy marker genes. The tree was visualized using iTOL [20].

### Structure prediction and alignment of PBP

Protein sequences were retrieved from the annotated genome files and structures were predicted with the AlphaFold server (AlphaFold3, accessed on 12th October 2026) [21] with a single protein chain as the only entity, default parameters and seed set to auto. Of the five models generated per sequence, the top-ranked model was used, protein sequence length and pTM values for every model are given in Supplementary Table 15. Pairwise structure alignments were performed with the RCSB PDBank Structure Alignment service using the jCE-CP algorithm with default parameters[22]. Full-length PBPs were aligned onto the PBP of HlEnr_12, truncated PBPs onto that of '*Ca.* Uabimicrobium amorphum' SRT547, and truncated PBPs onto the full-length protein of HlEnr_12. TM scores for all pairs are given in Supplementary Table 15. Superimposed structures were rendered in ChimeraX (version 1.11.1) [23]. For the comparison of full-length and truncated PBPs, the N-terminal α-helix of the truncated models (residue 1 to 48 or 49, depending on the model) was hidden from display to improve visibility of the shared core, superposition and reported statistics were calculated on the complete models. Pfam domains were predicted using InterProScan6 [24].

**BLASTp based horizontal gene transfer assessment M&M**

Genes were predicted for all genomes using prodigal [25], after which the resulting protein-coding sequences were queried against FunTaxDB 1.4 [26] with DIAMOND BLASTp [27] under --fast mode, -e 0.00001 and -k 1 to obtain only the first hit for each query and ensuring the minimum p-value is $10^{-5}$, so as to ascertain their closest function and taxonomy. Gene-level BLASTp results for each genome were imported into R using tidyverse (https://tidyverse.tidyverse.org/articles/paper.html), and their FunTaxDB taxonomy string parsed to obtain domain-level taxonomy (Bacteria, Archaea, Viruses) and to distinguish between *Planctomycetota* and other bacterial phyla. A barplot was generated using ggplot2 [28].

### 3. Supplementary Figures

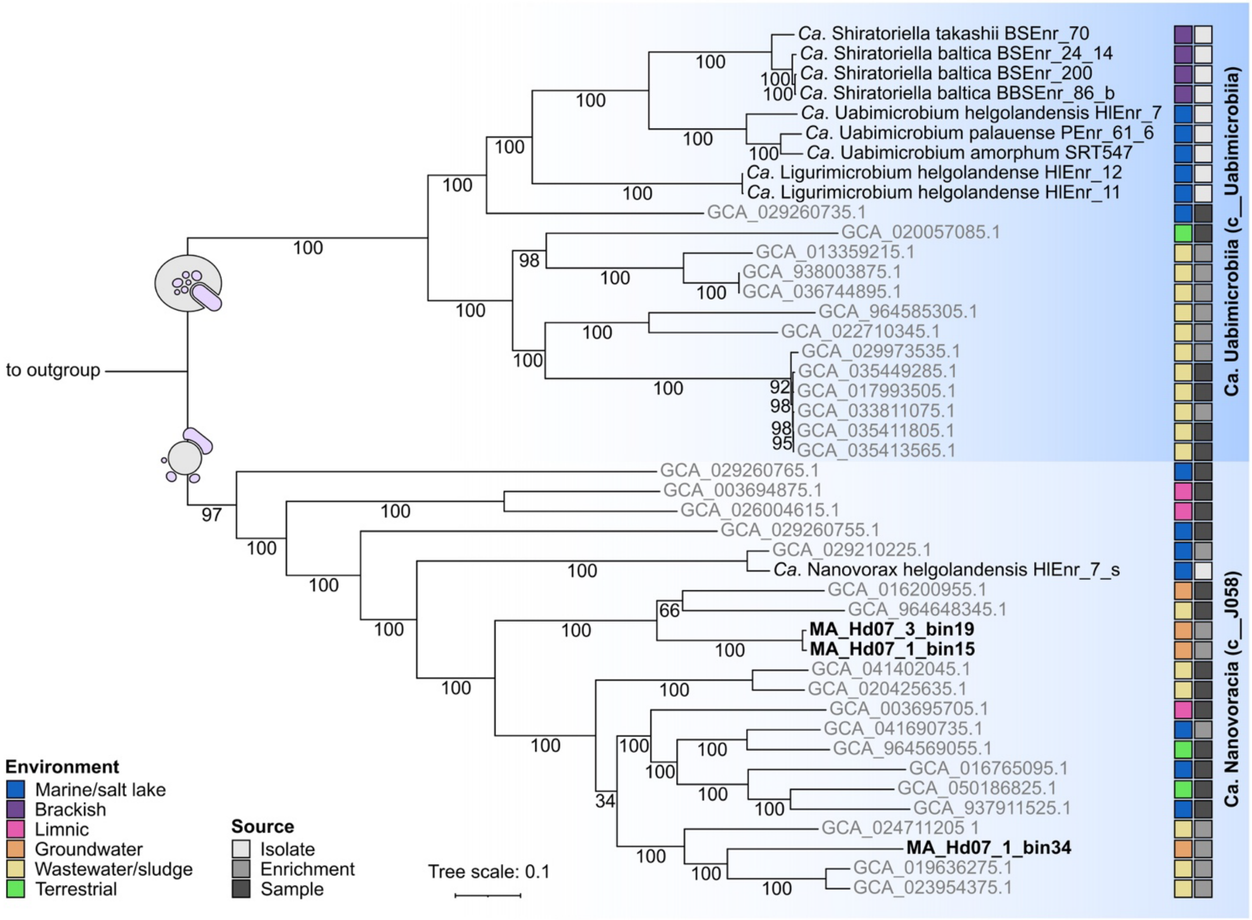


**Supplementary Fig. 1: Phylogenomic placement of cultured and metagenomically sequenced predatory *Planctomycetota*.**

Phylogenomic tree of the strains described in this study and related genomes, based on 71 concatenated bacterial single-copy marker genes. MAGs accessible from GTDB are displayed in grey font, isolates are displayed in black font and MAGs described in this study are highlighted by bold font. Environmental information is provided in the first column of squares; the source type of the genomic sequence is indicated in the second column. All nine bacteria of prey form a single clade together with the 13 other MAGs found in GTDB class c_Uabimicrobiia. The non-engulfing strain '*Ca.* Nanovorax helgolandensis' HlEnr_7_s clusters together with the MAGs obtained from groundwater enrichments and the 18 further MAGs found in GTDB class c__J058. Scale bar, 0.1 substitutions per site.

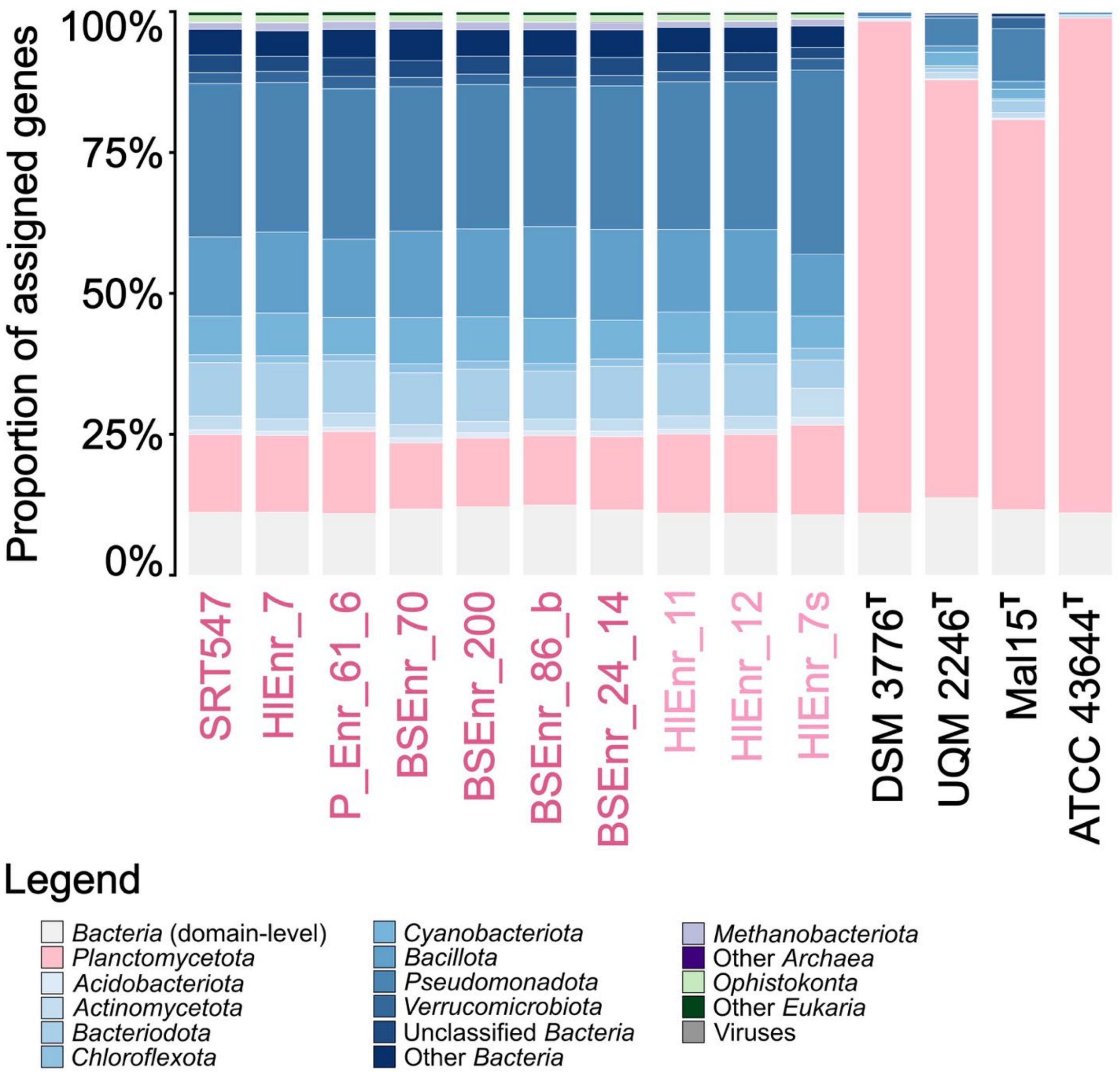


**Supplementary Fig. 2: Horizontal Gene Transfer in predatory *Planctomycetota*.**

Taxonomic context for genes found in predatory *Planctomycetota* in comparison to *Planctopirus limnophila* DSM3776$^{T}$, *Gemmata obscuriglobus* UQM 2246$^{T}$, *Stieleria maiorica* Mal15$^{T}$ and *Isosphaera pallida* ATCC 43644$^{T}$. In case of predatory *Planctomycetota* members, the fraction of genes with orthologs found in the same phylum is much lower than in case of the canonical model species, indicating high levels of horizontal gene transfer including even orthologs from other domains.

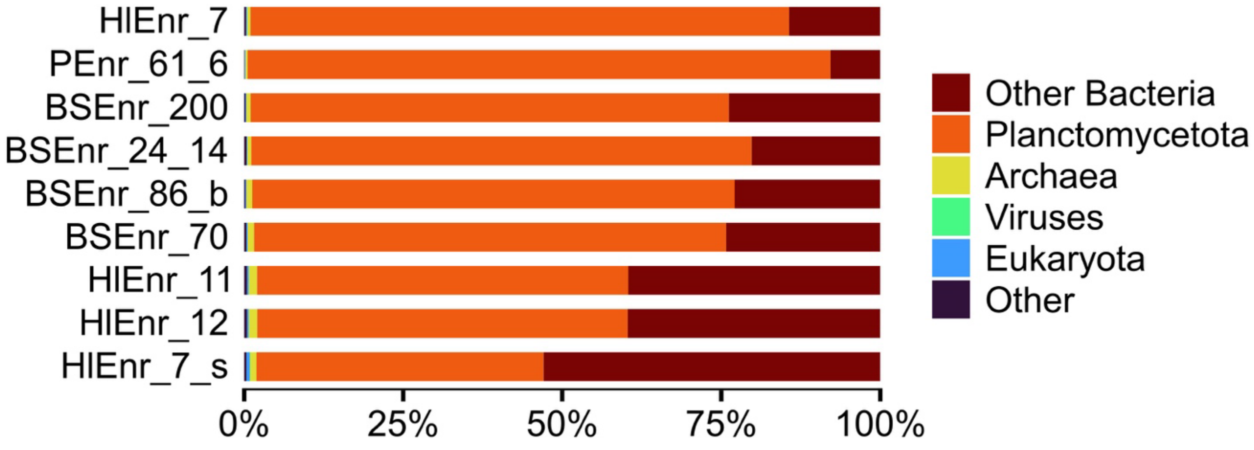


**Supplementary Fig. 3: Taxonomic affiliation of best BLASTp hits for the predicted proteomes of the nine novel strains.**

Each predicted protein of the eight phagocytic strains and the non-engulfing strain HlEnr_7_s was searched with BLASTp against FunTaxDB 1.4 [26], and the taxonomic origin of the best hit was recorded (e-value $\leq 10^{-5}$; proteins without a hit were excluded). Bars show the fraction of proteins whose best hit belongs to *Planctomycetota*, to other bacteria, to Archaea, Eukaryota, viruses, or to other taxa. In all nine genomes, 90–95 % or more of the proteins have best hits in *Planctomycetota* or other bacteria, with marginal contributions from archaea, eukaryotes and viruses. Strain HlEnr_7_s ('*Ca.* Nanovorax helgolandensis') shows the largest proportion of non-planctomycetal bacterial best hits. Note that a best hit indicates the taxonomic affiliation of the closest detectable homologue and does not by itself demonstrate horizontal gene transfer.

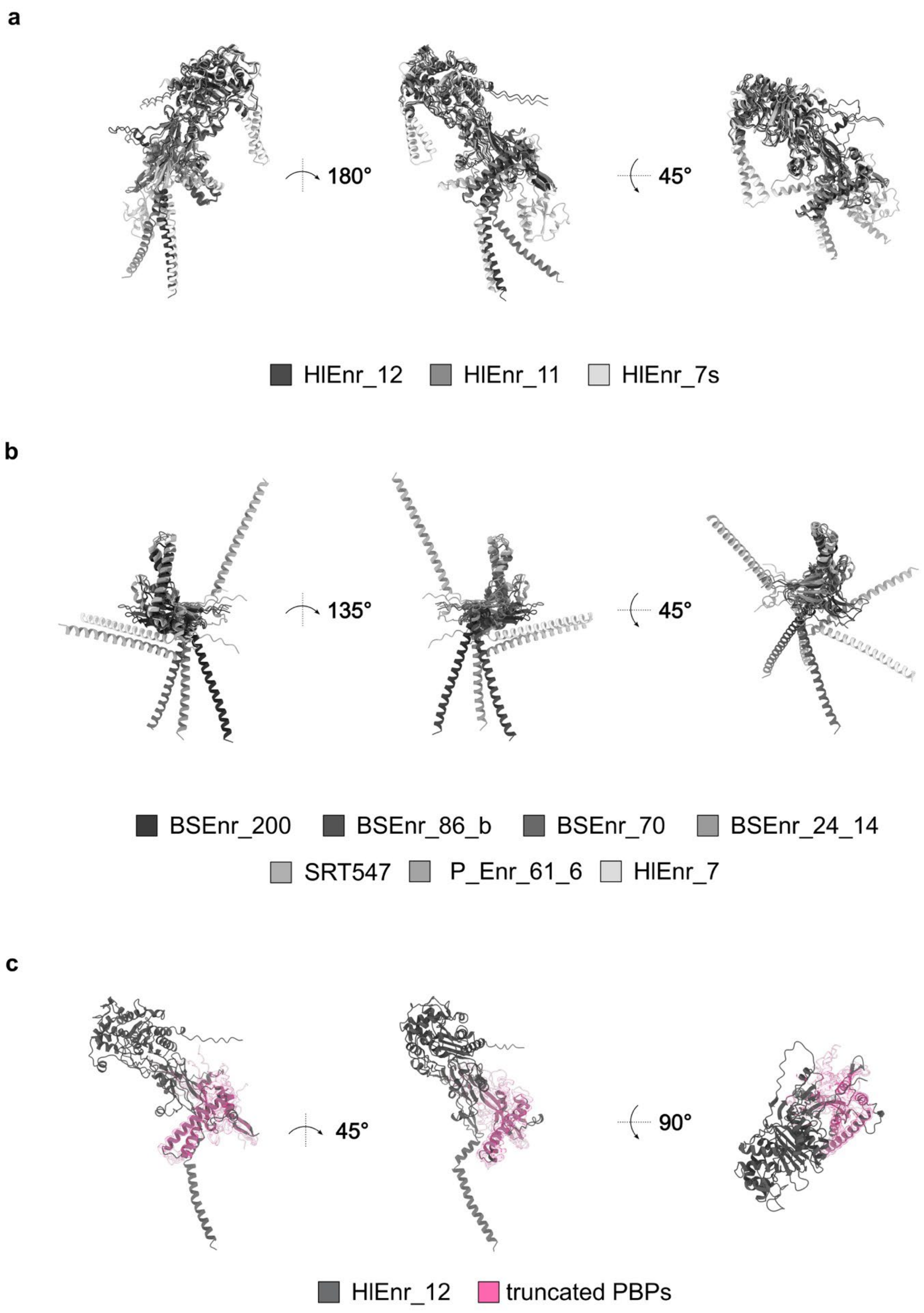
a
180°
45°
HlEnr_12
HlEnr_11
HlEnr_7s
b
135°
45°
BSEnr_200
BSEnr_86_b
BSEnr_70
BSEnr_24_14
SRT547
P_Enr_61_6
HlEnr_7
c
45°
90°
HlEnr_12
truncated PBPs

**Supplementary Fig. 4: Protein structure prediction and alignments of penicillin-binding proteins.**

(a) Full-length penicillin-binding proteins (PBP) could be identified in all strains dividing via cell division mode II (CDM II). While all three full-length PBPs share most of their domains, strain HlEnr_7s possess additional motifs. (b,c) In BoP dividing via CDM I, C-terminally truncated PBPs could be identified. (b) The truncated versions share a common motif, but the N-terminal alpha helix is not present in all of the truncated PBPs.(c) When superimposed onto PBP of strain HlEnr_12, the N-terminal domain appeared conserved (pink), suggesting a truncation of the larger C-terminal domains. N-terminal alpha helices of truncated versions were hidden in the superimposition for better visualization of the conserved domain illustrated in pink.

### Supplementary Tables

**Supplementary Table 1: Similarity matrix based on 16S rRNA gene sequences (%).** Thresholds used: species: 98.7%, genus: 94.5%

| | BSEnr_24_14 | BSEnr_200 and BSEnr_86_b | BSEnr_70 | PEnr_61_6 | SRT547 | HlEnr_7 | HlEnr_11 and HlEnr_12 | HlEnr_7_s |
|---|---|---|---|---|---|---|---|---|
| BSEnr_24_14 | 100.0 | 99.7 | 98.4 | 90.2 | 90.5 | 90.4 | 88.5 | 76.3 |
| BSEnr_200 and BSEnr_86_b | 99.7 | 100.0 | 98.5 | 90.2 | 90.4 | 90.4 | 88.5 | 76.2 |
| BSEnr_70 | 98.4 | 98.5 | 100.0 | 90.4 | 90.7 | 90.3 | 88.7 | 76.3 |
| PEnr_61_6 | 90.2 | 90.2 | 90.4 | 100.0 | 98.4 | 97.0 | 89.1 | 75.8 |
| SRT547 | 90.5 | 90.4 | 90.7 | 98.4 | 100.0 | 97.0 | 89.0 | 76.3 |
| HlEnr_7 | 90.4 | 90.4 | 90.3 | 97.0 | 97.0 | 100.0 | 88.8 | 76.3 |
| HlEnr_11 and HlEnr_12 | 88.5 | 88.5 | 88.7 | 89.1 | 89.0 | 88.8 | 100.0 | 76.8 |
| HlEnr_7_s | 76.3 | 76.2 | 76.3 | 75.8 | 76.3 | 76.3 | 76.8 | 100.0 |

**Supplementary Table 2: Similarity matrix based on average nucleotide identity values (ANI) (%).** Thresholds used: species: 95%, no commonly used genus threshold.

| | BSEnr_200 | BSEnr_24_14 | BSEnr_70 | BSEnr_86_b | HlEnr_11 | HlEnr_12 | HlEnr_7 | HlEnr_7_s | PEnr_61_6 | SRT547 |
|---|---|---|---|---|---|---|---|---|---|---|
| BSEnr_200 | 100.0 | 96.3 | 84.4 | 98.7 | 77.7 | 78.1 | 74.8 | < 70.0 | 75.3 | 74.7 |
| BSEnr_24_14 | 96.3 | 100.0 | 85.0 | 96.6 | 78.6 | 78.6 | 74.7 | < 70.0 | 75.7 | 74.3 |
| BSEnr_70 | 84.4 | 85.0 | 100.0 | 84.7 | 76.4 | 77.0 | 74.6 | < 70.0 | 76.6 | 74.7 |
| BSEnr_86_b | 98.7 | 96.6 | 84.7 | 100.0 | 77.0 | 76.9 | 75.5 | < 70.0 | 75.6 | 74.9 |
| HlEnr_11 | 77.7 | 78.6 | 76.4 | 77.0 | 100.0 | 100.0 | 80.3 | < 70.0 | 78.9 | 80.4 |
| HlEnr_12 | 78.1 | 78.6 | 77.0 | 76.9 | 100.0 | 100.0 | 80.5 | < 70.0 | 78.7 | 80.2 |
| HlEnr_7 | 74.8 | 74.7 | 74.6 | 75.5 | 80.3 | 80.5 | 100.0 | < 70.0 | 77.9 | 77.6 |
| HlEnr_7_s | < 70.0 | < 70.0 | < 70.0 | < 70.0 | < 70.0 | < 70.0 | < 70.0 | 100.0 | < 70.0 | < 70.0 |
| PEnr_61_6 | 75.3 | 75.7 | 76.6 | 75.6 | 78.9 | 78.7 | 77.9 | < 70.0 | 100.0 | 81.2 |
| SRT547 | 74.7 | 74.3 | 74.7 | 74.9 | 80.4 | 80.2 | 77.6 | < 70.0 | 81.2 | 100.0 |

**Supplementary Table 3: Similarity matrix based on average amino acid identity values (AAI) (%).** Thresholds used: species: 95%, genus: 60-80%.

| | BSEnr_200 | BSEnr_24_14 | BSEnr_70 | BSEnr_86_b | HIEnr_11 | HIEnr_12 | HIEnr_7 | HIEnr_7_s | PEnr_61_6 | SRT547 |
|---|---|---|---|---|---|---|---|---|---|---|
| BSEnr_200 | 100.0 | 96.3 | 86.7 | 98.2 | 49.3 | 49.3 | 57.6 | 36.9 | 57.7 | 57.5 |
| BSEnr_24_14 | 96.3 | 100.0 | 86.8 | 96.6 | 49.4 | 49.4 | 57.8 | 36.9 | 57.8 | 57.5 |
| BSEnr_70 | 86.7 | 86.8 | 100.0 | 86.7 | 49.2 | 49.2 | 57.6 | 36.7 | 57.8 | 57.2 |
| BSEnr_86_b | 98.2 | 96.6 | 86.7 | 100.0 | 49.3 | 49.3 | 57.8 | 36.7 | 57.8 | 57.6 |
| HIEnr_11 | 49.3 | 49.4 | 49.2 | 49.3 | 100.0 | 100.0 | 49.5 | 37.3 | 49.6 | 49.5 |
| HIEnr_12 | 49.3 | 49.4 | 49.2 | 49.3 | 100.0 | 100.0 | 49.5 | 37.3 | 49.6 | 49.5 |
| HIEnr_7 | 57.6 | 57.8 | 57.6 | 57.8 | 49.5 | 49.5 | 100.0 | 36.9 | 72.7 | 71.7 |
| HIEnr_7_s | 36.9 | 36.9 | 36.7 | 36.7 | 37.3 | 37.3 | 36.9 | 100.0 | 37.1 | 36.9 |
| PEnr_61_6 | 57.7 | 57.8 | 57.8 | 57.8 | 49.6 | 49.6 | 72.7 | 37.1 | 100.0 | 83.7 |
| SRT547 | 57.5 | 57.5 | 57.2 | 57.6 | 49.5 | 49.5 | 71.7 | 36.9 | 83.7 | 100.0 |

**Supplementary Table 4: Similarity matrix based on partial *rpoB* gene similarity (%).** Thresholds used: species: 96.3%, genus: 75.5-78.0%.

| | PEnr_61_6 | SRT547 | HlEnr_7 | BSEnr_200 | BSEnr_86_b | BSEnr_24_14 | BSEnr_70 | HlEnr_11 and HlEnr_12 | HlEn7_s |
|---|---|---|---|---|---|---|---|---|---|
| PEnr_61_6 | 100.0 | 88.7 | 81.7 | 73.5 | 73.5 | 73.2 | 72.3 | 69.0 | 56.8 |
| SRT547 | 88.7 | 100.0 | 82.8 | 73.5 | 73.6 | 73.3 | 72.8 | 69.6 | 57.5 |
| HlEnr_7 | 81.7 | 82.8 | 100.0 | 75.3 | 75.5 | 75.7 | 75.4 | 69.4 | 58.1 |
| BSEnr_200 | 73.5 | 73.5 | 75.3 | 100.0 | 99.4 | 98.1 | 88.0 | 70.2 | 56.6 |
| BSEnr_86_b | 73.5 | 73.6 | 75.5 | 99.4 | 100.0 | 98.1 | 88.2 | 70.3 | 56.5 |
| BSEnr_24_14 | 73.2 | 73.3 | 75.7 | 98.1 | 98.1 | 100.0 | 88.9 | 70.2 | 56.3 |
| BSEnr_70 | 72.3 | 72.8 | 75.4 | 88.0 | 88.2 | 88.9 | 100.0 | 69.5 | 56.6 |
| HlEnr_11 and HlEnr_12 | 69.0 | 69.6 | 69.4 | 70.2 | 70.3 | 70.2 | 69.5 | 100.0 | 57.8 |
| HlEn7_s | 56.8 | 57.5 | 58.1 | 56.6 | 56.5 | 56.3 | 56.6 | 57.8 | 100.0 |

**Supplementary Table 5: Similarity matrix based on values for percentage of conserved proteins (POCP) (%).** Thresholds: species: no commonly used threshold, genus: 50%.

| | BSEnr_200 | BSEnr_24_14 | BSEnr_70 | BSEnr_86_b | HlEnr_11 | HlEnr_12 | HlEnr_7 | HlEnr_7_s | PEnr_61_6 | SRT547 |
|---|---|---|---|---|---|---|---|---|---|---|
| BSEnr_200 | 100.0 | 86.8 | 80.1 | 89.7 | 41.0 | 41.0 | 56.4 | 11.7 | 59.4 | 55.5 |
| BSEnr_24_14 | 86.8 | 100.0 | 81.6 | 87.4 | 41.5 | 41.5 | 57.7 | 11.8 | 60.6 | 56.6 |
| BSEnr_70 | 80.1 | 81.6 | 100.0 | 81.1 | 40.7 | 40.7 | 56.2 | 11.1 | 58.2 | 54.3 |
| BSEnr_86_b | 89.7 | 87.4 | 81.1 | 100.0 | 41.2 | 41.2 | 56.7 | 11.6 | 60.1 | 56.1 |
| HlEnr_11 | 41.0 | 41.5 | 40.7 | 41.2 | 100.0 | 99.3 | 42.1 | 12.0 | 42.8 | 41.6 |
| HlEnr_12 | 41.0 | 41.5 | 40.7 | 41.2 | 99.3 | 100.0 | 42.0 | 12.0 | 42.8 | 41.5 |
| HlEnr_7 | 56.4 | 57.7 | 56.2 | 56.7 | 42.1 | 42.0 | 100.0 | 11.4 | 72.7 | 70.0 |
| HlEnr_7_s | 11.7 | 11.8 | 11.1 | 11.6 | 12.0 | 12.0 | 11.4 | 100.0 | 12.2 | 11.9 |
| PEnr_61_6 | 59.4 | 60.6 | 58.2 | 60.1 | 42.8 | 42.8 | 72.7 | 12.2 | 100.0 | 79.0 |
| SRT547 | 55.5 | 56.6 | 54.3 | 56.1 | 41.6 | 41.5 | 70.0 | 11.9 | 79.0 | 100.0 |

**Supplementary Table 6: Overview on genomic features of novel bacteria of prey.** Hypothetical protein counts are based on the PGAP / RefSeq annotation.

| Characteristics | SRT547 | HIEnr_7 | PEnr_61_6 | BSEnr_70 | BSEnr_24_14 | BSEnr_200 | BSEnr_86_b | HIEnr_11 | HIEnr_12 | HIEnr_7_s |
|---|---|---|---|---|---|---|---|---|---|---|
| Genome size (bp) | 9,503,417 | 9,369,075 | 7,958,823 | 7,395,093 | 6,959,158 | 7,166,336 | 7,052,067 | 9,854,279 | 9,875,819 | 10,372,370 |
| Contigs | 1 | 1 | 1 | 2 | 1 | 2 | 1 | 3 | 4 | 1 |
| Plasmids | 0 | 0 | 0 | 1 | 0 | 1 | 0 | 2 | 3 | 0 |
| DNA G+C content (%) | 39.4 | 34.9 | 38.9 | 34.4 | 34.3 | 34.3 | 34.3 | 35.5 | 35.4 | 51.5 |
| Genes | 6,712 | 6,933 | 5,871 | 5,657 | 5,153 | 5,237 | 5,105 | 7,021 | 7,044 | 7,633 |
| Genes/Mbp | 706 | 740 | 738 | 765 | 740 | 731 | 724 | 712 | 713 | 736 |
| Protein-coding genes | 6,585 | 6,725 | 5,740 | 5,526 | 5,055 | 5,130 | 4,984 | 6,868 | 6,894 | 7,484 |
| Protein-coding genes / Mbp | 693 | 718 | 721 | 747 | 726 | 716 | 707 | 697 | 698 | 722 |
| Hypothetical proteins | 1,705 | 1,740 | 1,348 | 1,466 | 1,248 | 1,213 | 1,140 | 1,760 | 1,774 | 2,081 |
| Hypothetical proteins (%) | 25.9 | 25.9 | 23.5 | 26.5 | 24.7 | 23.6 | 22.9 | 25.6 | 25.7 | 27.8 |
| Coding density (%) | 91.4 | 87.4 | 91.1 | 88.7 | 88.8 | 89.8 | 89.6 | 89.0 | 89.0 | 90.5 |
| rRNA genes (5S,16S,23S) | 3,3,3 | 4,4,4 | 4,4,4 | 4,4,4 | 4,3,4 | 3,3,3 | 4,3,3 | 3,3,3 | 3,3,3 | 4,4,4 |
| tRNA genes | 76 | 88 | 67 | 44 | 45 | 45 | 46 | 82 | 84 | 88 |

**Supplementary Table 7: Analysis of secondary metabolite-associated biosynthetic gene clusters (BGCs) using antiSMASH.** PKS: polyketide synthase, NRPS: non-ribosomal peptide synthetase, RiPP: ribosomally synthesized and posttranslationally modified peptide, RRE: RiPP recognition element

| Characteristics | SRT547 | HIEnr_7 | PEnr_61_6 | BSEnr_70 | BSEnr_24_14 | BSEnr_200 | BSEnr_86_b | HIEnr_11 | HIEnr_12 | HIEnr_7_s |
|---|---|---|---|---|---|---|---|---|---|---|
| Terpene (incl. precursors) | 8 | 7 | 8 | 4 | 3 | 4 | 4 | 4 | 4 | 2 |
| Type I PKS | 2 | 3 | 2 | 5 | 7 | 8 | 8 | 11 | 11 | 2 |
| Type II PKS | 1 | 1 | 1 | 1 | 1 | 1 | 1 | 0 | 0 | 0 |
| Type III PKS | 1 | 1 | 1 | 1 | 1 | 1 | 1 | 1 | 1 | 1 |
| NRPS | 3 | 4 | 4 | 2 | 2 | 2 | 1 | 2 | 2 | 0 |
| NRPS-like | 3 | 3 | 2 | 4 | 4 | 1 | 5 | 6 | 6 | 0 |
| Mixed type I PKS-NRPS | 3 | 3 | 2 | 5 | 4 | 3 | 3 | 1 | 1 | 0 |
| Oligosaccharide | 1 | 0 | 0 | 0 | 0 | 0 | 0 | 0 | 0 | 0 |
| RiPP/RiPP-like | 2 | 2 | 2 | 2 | 3 | 3 | 2 | 3 | 3 | 6 |
| Arylpolyene | 2 | 2 | 1 | 2 | 2 | 2 | 2 | 0 | 0 | 0 |
| RRE-containing | 0 | 1 | 1 | 2 | 0 | 0 | 0 | 2 | 2 | 0 |
| Lassopeptide | 0 | 0 | 1 | 0 | 0 | 0 | 0 | 0 | 0 | 0 |
| Lanthipeptide | 0 | 0 | 0 | 0 | 1 | 0 | 0 | 1 | 1 | 0 |
| Phosphonate | 0 | 0 | 0 | 0 | 0 | 0 | 0 | 1 | 1 | 0 |
| Total | 26 | 27 | 25 | 28 | 28 | 25 | 27 | 32 | 32 | 11 |

**Supplementary Table 8: Genome-based analysis of carbohydrate-active enzymes (CAZymes).** The run_dbCAN tool was applied.

| Characteristics | SRT547 | HIEnr_7 | PEnr_61_6 | BSEnr_70 | BSEnr_24_14 | BSEnr_200 | BSEnr_86_b | HIEnr_11 | HIEnr_12 | HIEnr_7_s | *Planctopirus limnophila* DSM 3776 | *Stieleria maiorica* Mal15 |
|---|---|---|---|---|---|---|---|---|---|---|---|---|
| Glycoside hydrolases | 24 | 17 | 24 | 14 | 13 | 13 | 12 | 29 | 29 | 24 | 32 | 81 |
| Glycosyltransferases | 82 | 61 | 65 | 71 | 54 | 61 | 60 | 79 | 79 | 58 | 60 | 102 |
| Polysaccharide lyases | 1 | 1 | 2 | 1 | 1 | 0 | 0 | 1 | 1 | 6 | 2 | 8 |
| Carbohydrate esterases | 6 | 4 | 5 | 3 | 3 | 5 | 4 | 5 | 5 | 4 | 15 | 28 |
| Carbohydrate-bind. modules | 5 | 7 | 5 | 5 | 5 | 5 | 5 | 7 | 7 | 4 | 14 | 21 |
| Auxiliary activities | 4 | 2 | 1 | 3 | 3 | 2 | 2 | 4 | 4 | 3 | 2 | 3 |
| Total | 122 | 92 | 102 | 97 | 79 | 86 | 83 | 125 | 125 | 99 | 125 | 243 |

**Supplementary Table 9: Curated list of 192 eukaryotic phagocytosis proteins.**

See supplementary file.

**Supplementary Table 10: Sequence-based identified homologous of eukaryotic phagocytosis proteins in predatory *Planctomycetota*.**

See supplementary file.

**Supplementary Table 11: Orthogroups exclusively found in bacteria of prey.**

See supplementary file.

**Supplementary Table 12: Orthogroups associated to specific cell division modes.**

See supplementary file.

**Supplementary Table 13: Sampling information, culture media, and accession numbers.** All samples were taken from surface water. D: Germany, P: Palau; sW: sterile filtered water from the sampling site; JCM: JCM Medium 1273 with artificial seawater from JCM Medium 1118 without LB; TrM: TrM35 artificial sea water

| Strain | sampling date | habitat | geographic location | latitude | longitude | culture medium | BioSample number | 16S accession number |
|---|---|---|---|---|---|---|---|---|
| HlEnr_7 | 22.06.23 | marine | Heligoland (D) | 54.1832656 | 7.8900549 | sW | SAMN38242342 | OR832863 |
| HlEnr_7_s | 22.06.23 | marine | Heligoland (D) | 54.1832656 | 7.8900549 | JCM | SAMN62694635 | PZ940871 |
| HlEnr_11 | 22.06.23 | marine | Heligoland (D) | 54.1832656 | 7.8900549 | sW | SAMN62694636 | PZ940872 |
| HlEnr_12 | 22.06.23 | marine | Heligoland (D) | 54.1832656 | 7.8900549 | sW | SAMN62694649 | PZ940873 |
| BSEnr_70 | 27.07.23 | brackish | Nienhagen (D) | 54.1648519 | 11.9463111 | sW | SAMN62694687 | PZ940876 |
| BSEnr_200 | 30.07.23 | brackish | Dierhagen (D) | 54.302472 | 12.34253 | sW | SAMN62694728 | PZ940877 |
| BSEnr_86_b | 27.07.23 | brackish | Prerow (D) | 54.4509496 | 12.6029807 | sW | SAMN62694759 | PZ940878 |
| BSEnr_24_14 | 21.09.24 | brackish | Graal-Müritz (D) | 54.253213 | 12.22086 | sW | SAMN62694832 | PZ940879 |
| PEnr_61_6 | 03.02.25 | marine | Wonder Lake (P) | 7.31025 | 134.494556 | TrM | SAMN62694837 | PZ947348 |

**Supplementary Table 14: Sequencing and genome assembly details.**

See supplementary file.

**Supplementary Table 15: Data from protein structure prediction (AlphaFold3) and alignment scores (RCSB PDBank Structure Alignment service, jCE-CP algorithm).**

| Strain | Length | pTM | Overlay full-length TM | Overlay truncated TM | Overlay both TM | PF03717 | PF00905 |
|---|---|---|---|---|---|---|---|
| SRT547 | 199 | 0.31 | | Reference | 0.3 | absent | absent |
| HlEnr_7 | 197 | 0.61 | | 0.33 | 0.5 | absent | absent |
| P_Enr_61_6 | 200 | 0.4 | | 0.33 | 0.42 | absent | absent |
| BSEnr_70 | 152 | 0.72 | | 0.37 | 0.72 | absent | absent |
| BSEnr_200 | 199 | 0.59 | | 0.33 | 0.58 | absent | absent |
| BSEnr_86b | 199 | 0.57 | | 0.33 | 0.58 | absent | absent |
| BSEnr_24_14 | 199 | 0.56 | | 0.37 | 0.56 | absent | absent |
| HlEnr_11 | 676 | 0.82 | 0.89 | | | 54-287 | 314-655 |
| HlEnr_12 | 676 | 0.81 | Reference | | Reference | 54-287 | 314-655 |
| HlEnr_7s | 783 | 0.79 | 0.61 | | | 218-294 | 424-774 |

### Supplementary References


1 Sharma, A., Küsel, K., Wegner, C. E., Pérez-Carrascal, O. M. & Taubert, M. Two worlds beneath: Distinct microbial strategies of the rock-attached and planktonic subsurface biosphere. *Microbiome* **14**, 73 (2026). https://doi.org:10.1186/s40168-025-02325-1

2 Sharma, A. *et al.* Iron coatings on carbonate rocks shape the attached bacterial aquifer community. *Sci Total Environ* **917**, 170384 (2024). https://doi.org:10.1016/j.scitotenv.2024.170384

3 Herrmann, M. *et al.* Large fractions of $CO_2$-fixing microorganisms in pristine limestone aquifers appear to be involved in the oxidation of reduced sulfur and nitrogen compounds. *Appl Environ Microb* **81**, 2384-2394 (2015). https://doi.org:10.1128/Aem.03269-14

4 Herrmann, M. *et al.* Predominance of *Cand.* Patescibacteria in groundwater is caused by their preferential mobilization from soils and flourishing under oligotrophic conditions. *Front Microbiol* **10**, 1407 (2019). https://doi.org:10.3389/Fmicb.2019.01407

5 Küsel, K. *et al.* How deep can surface signals be traced in the Critical Zone? Merging biodiversity with biogeochemistry research in a Central German Muschelkalk landscape. *Front Earth Sci* **4**, 1-18 (2016). https://doi.org:10.3389/Feart.2016.00032

6 Lehmann, R. & Totsche, K. U. Multi-directional flow dynamics shape groundwater quality in sloping bedrock strata. *J Hydrol* **580**, 124291 (2020). https://doi.org:10.1016/j.jhydrol.2019.124291

7 Bushnell, B. BBTools software package. *URL http://sourceforge. net/projects/bbmap* (2014).

8 Nurk, S., Meleshko, D., Korobeynikov, A. & Pevzner, P. A. metaSPAdes: a new versatile metagenomic assembler. *Genome Res* **27**, 824-834 (2017). https://doi.org:10.1101/gr.213959.116

9 Alneberg, J. *et al.* Binning metagenomic contigs by coverage and composition. *Nat Methods* **11**, 1144-1146 (2014). https://doi.org:10.1038/Nmeth.3103

10 Kang, D. W. D. *et al.* MetaBAT 2: an adaptive binning algorithm for robust and efficient genome reconstruction from metagenome assemblies. *Peerj* **7**, 1-13 (2019). https://doi.org:10.7717/Peerj.7359

11 Wu, Y. W., Simmons, B. A. & Singer, S. W. MaxBin 2.0: an automated binning algorithm to recover genomes from multiple metagenomic datasets. *Bioinformatics* **32**, 605-607 (2016). https://doi.org:10.1093/bioinformatics/btv638

12 Uritskiy, G. V., DiRuggiero, J. & Taylor, J. MetaWRAP - a flexible pipeline for genome-resolved metagenomic data analysis. *Microbiome* **6**, 1-13 (2018). https://doi.org:10.1186/S40168-018-0541-1

13 Graham, E. D., Heidelberg, J. F. & Tully, B. J. BinSanity: unsupervised clustering of environmental microbial assemblies using coverage and affinity propagation. *Peerj* **5**, 1-19 (2017). https://doi.org:10.7717/Peerj.3035

14 Brown, C. T. *et al.* Unusual biology across a group comprising more than 15% of domain Bacteria. *Nature* **523**, 208-173 (2015). https://doi.org:10.1038/nature14486

15 Olm, M. R., Brown, C. T., Brooks, B. & Banfield, J. F. dRep: a tool for fast and accurate genomic comparisons that enables improved genome recovery from metagenomes through de-replication. *ISME J* **11**, 2864-2868 (2017). https://doi.org:10.1038/ismej.2017.126

16 Parks, D. H., Imelfort, M., Skennerton, C. T., Hugenholtz, P. & Tyson, G. W. CheckM: assessing the quality of microbial genomes recovered from isolates, single cells, and metagenomes. *Genome Res* **25**, 1043-1055 (2015). https://doi.org:10.1101/gr.186072.114

17 Chaumeil, P. A., Mussig, A. J., Hugenholtz, P. & Parks, D. H. GTDB-Tk: a toolkit to classify genomes with the Genome Taxonomy Database. *Bioinformatics* **36**, 1925-1927 (2019). https://doi.org:10.1093/bioinformatics/btz848

18 Aroney, S. T. N. *et al.* CoverM: read alignment statistics for metagenomics. *Bioinformatics* **41**, btaf147 (2025). https://doi.org:10.1093/bioinformatics/btaf147

19 Eren, A. M. *et al.* Community-led, integrated, reproducible multi-omics with anvi'o. *Nat Microbiol* **6**, 3-6 (2021). https://doi.org:10.1038/s41564-020-00834-3

20 Letunic, I. & Bork, P. Interactive Tree of Life (iTOL) v6: recent updates to the phylogenetic tree display and annotation tool. *Nucleic acids research* **52**, W78-W82 (2024). https://doi.org:10.1093/nar/gkae268

21 Abramson, J. *et al.* Accurate structure prediction of biomolecular interactions with AlphaFold 3. *Nature* **630**, 493-500 (2024). https://doi.org:10.1038/s41586-024-07487-w

22 Bittrich, S., Segura, J., Duarte, J. M., Burley, S. K. & Rose, Y. RCSB protein Data Bank: exploring protein 3D similarities via comprehensive structural alignments. *Bioinformatics* **40**, btae370 (2024). https://doi.org:10.1093/bioinformatics/btae370

23 Pettersen, E. F. *et al.* UCSF ChimeraX: Structure visualization for researchers, educators, and developers. *Protein Science* **30**, 70-82 (2021). https://doi.org:https://doi.org/10.1002/pro.3943

24 Blum, M., Hobbs, E., Florentino, L. & Bateman, A. InterProScan 6: a modern large-scale protein function annotation pipeline. *Bioinformatics Advances* **6**, vbag141 (2026). https://doi.org:10.1093/bioadv/vbag141

25 Hyatt, D. *et al.* Prodigal: prokaryotic gene recognition and translation initiation site identification. *BMC bioinformatics* **11** (2010). https://doi.org:10.1186/1471-2105-11-119

26 Bornemann, T. L. V., Esser, S. P., Stach, T. L., Burg, T. & Probst, A. J. uBin: A manual refining tool for genomes from metagenomes. *Environmental Microbiology* **25**, 1077-1083 (2023). https://doi.org:https://doi.org/10.1111/1462-2920.16351

27 Buchfink, B., Xie, C. & Huson, D. H. Fast and sensitive protein alignment using DIAMOND. *Nature methods* (2014). https://doi.org:10.1038/nmeth.3176

28 Wickham, H. *ggplot2: elegant graphics for data analysis*. (Springer New York, 2009).
